# Diets affect the age-dependent decline of associative learning in *Caenorhabditis elegans*

**DOI:** 10.1101/2020.10.08.331256

**Authors:** Satoshi Higurashi, Sachio Tsukada, Binta Maria Aleogho, Joo Hyun Park, Masaru Tanaka, Shunji Nakano, Ikue Mori, Kentaro Noma

**Author notes:** **Correspondence to** Kentaro Noma. Group of Microbial Motility, Department of Biological Science, Division of Natural Science, Graduate school of Science, Nagoya University, Nagoya, 464-8602, Japan (Kentaro Noma). These authors equally contributed to this work (Satoshi Higurashi, Sachio Tsukada).

## Abstract

The causality and mechanism of dietary effects on brain aging are still unclear due to the long time scales of aging. The nematode *Caenorhabditis elegans* has contributed to aging research because of its short lifespan and easy genetic manipulation. When fed the standard laboratory diet, *Escherichia coli*, *C. elegans* experiences an age-dependent decline in temperature-food associative learning, called thermotaxis. To address if diet affects this decline, we screened 35 lactic acid bacteria as alternative diets and found that animals maintained high thermotaxis ability when fed a clade of *Lactobacilli* enriched with heterofermentative bacteria. Among them, *Lactobacill*us *reuteri* maintained the thermotaxis of aged animals without affecting their lifespan and motility. The effect of *Lb. reuteri* depends on the DAF-16 transcription factor functioning in neurons. Furthermore, RNA sequencing analysis revealed that differentially expressed genes between aged animals fed different bacteria were enriched with DAF-16 targets. Our results demonstrate that diet can impact brain aging in a *daf-16* dependent manner without changing the lifespan.

## Introduction

Human life expectancy has increased since the nineteenth century (Dong, Milholland, & Vijg, 2016), which has led to the social problem related to age-dependent cognitive dysfunction. Although human studies suggest that genetic background, diet, and lifestyle might affect brain aging, the causality and mechanism of how they affect brain aging remain unclear (Deary et al., 2009).

The nematode *Caenorhabditis elegans* (*C. elegans*) is ideal for addressing the mechanism of age-related phenotypes because of the two to three-week lifespan and the variety of available genetic tools. In *C. elegans*, the age-related phenotypes can be readily separable from the organismal lifespan by directly measuring the lifespan. In the past decades, studies using *C. elegans* have contributed to aging research by revealing the molecular mechanism of how dietary restriction, insulin-like signaling, and germline stem cells affect organismal lifespan (Mack, Heimbucher, & Murphy, 2018; Wolff & Dillin, 2006). Like mammals, *C. elegans* experiences age-dependent functional changes in the nervous system (Stein & Murphy, 2012). Aged animals are defective in locomotion (Hahm et al., 2015; Mulcahy, Holden-Dye, & O’Connor, 2013), mechanosensory response (Beck & Rankin, 1993), chemotaxis (Leinwand et al., 2015), thermotaxis (Huang et al., 2020; H. Murakami, Bessinger, Hellmann, & Murakami, 2005; S. Murakami & Murakami, 2005), and food-butanone associative learning (Kauffman, Ashraf, Corces-Zimmerman, Landis, & Murphy, 2010). Age-dependent memory decline in the food-butanone association is ameliorated in the mutant of *nkat-1* encoding kynurenic acid-synthesizing enzyme prevents (Vohra, Lemieux, Lin, & Ashrafi, 2018). Overactivation of Gα signaling in AWC sensory neurons also maintains the ability to form memory in aged animals in the food-butanone association (Arey, Stein, Kaletsky, Kauffman, & Murphy, 2018). These emerging evidence suggests that genetic manipulations can prevent age-dependent functional decline in the nervous system.

Compared to genetic manipulations, the modification of diet can be easily applicable to our daily lives. Studies in humans and mice imply that diets affect the cognitive decline in aged animals (Joseph, Cole, Head, & Ingram, 2009; Vauzour et al., 2017). Here, we use *C. elegans* to address the dietary effect on the age-dependent behavioral decline and its underlying mechanism. In laboratories, *C. elegans* is maintained monoxenically with a uracil auxotroph *Escherichia coli* (*E. coli*) strain, OP50, as the standard diet (Brenner, 1974). On the other hand, *C. elegans* in natural habitat eats a wide variety of bacteria (Berg et al., 2016; Dirksen et al., 2016; Johnke, Dirksen, & Schulenburg, 2020; Samuel, Rowedder, Braendle, Felix, & Ruvkun, 2016; Zhang et al., 2017). These bacteria affect the physiology of *C. elegans*, such as growth rate, reproduction, and sensory behavior (Dirksen et al., 2016; O’Donnell, Fox, Chao, Schroeder, & Sengupta, 2020; Samuel et al., 2016). However, the effect of different bacteria on the behavioral decline during aging is unexplored. Among the potential bacterial diet for *C. elegans* in natural habitat (Berg et al., 2016; Dirksen et al., 2016; Samuel et al., 2016), we focused on Lactic Acid Bacteria (LAB), which are the most common probiotics for humans (Hill et al., 2014). LAB, such as *Lactobacilli* (*Lb.*) and *Bifidobacteria* (*B.*), are Gram-positive, non-spore-forming bacteria that produce lactic acid from carbohydrates as the primary metabolic product. Depending on the species, LAB have various effects on *C. elegans* physiology. *Lb. gasseri*, *B. longum*, and *B. infantis* extend lifespan in *C. elegans* (Komura, Ikeda, Yasui, Saeki, & Nishikawa, 2013; Nakagawa et al., 2016; L. Zhao et al., 2017). On the other hand, *Lb. helveticus* does not increase the lifespan (Nakagawa et al., 2016). Even in the same species, different strains have different effects on lifespan, body size, and locomotion (Wang et al., 2020). In *C. elegans*, LAB modulate evolutionarily conserved genetic pathways such as the insulin/insulin-like growth factor-1 (IGF-1) signaling (IIS) pathway (Grompone et al., 2012; Sugawara & Sakamoto, 2018), which consists of the insulin receptor DAF-2, phosphoinositide 3 (PI3) kinase cascade, and the downstream transcription factor DAF-16 (Kenyon, Chang, Gensch, Rudner, & Tabtiang, 1993; Lin, Dorman, Rodan, & Kenyon, 1997). DAF-16 is a sole *C. elegans* ortholog of mammalian FOXO transcription factor and is involved in various biological processes (Stein & Murphy, 2012; Tissenbaum, 2018).

To comprehensively understand the effect of LAB, we screened 35 different LAB species, including some subspecies. We examined the age-dependent functional decline of thermotaxis behavior, which reflects associative learning between temperature and food (Hedgecock & Russell, 1975; Mori & Ohshima, 1995). We demonstrate that *C. elegans* fed *Lactobacilli* in a clade maintained the thermotaxis behavior when aged, while *E. coli*-fed animals did not. Among those *Lactobacilli*, *Lb. reuteri* maintained the thermotaxis ability of aged animals without affecting the organismal lifespan or locomotion. The effect of *Lb. reuteri* on the thermotaxis of aged animals depends on the DAF-16 transcription factor functioning in neurons.

## Results

### *C. elegans* thermotaxis behavior declines with age

After being cultivated with food at a temperature within the physiological range (15∼25 °C), *C. elegans* migrates toward and stays at the past cultivation temperature (T_cult_) on a linear thermal gradient without food (Fig. 1A). This behavior is called thermotaxis (Hedgecock & Russell, 1975; Mori & Ohshima, 1995). To see the effect of aging on thermotaxis, we cultivated animals at 20 °C with an *E. coli* strain, OP50 (hereafter, *E. coli*, unless otherwise noted), commonly used in laboratory conditions (Brenner, 1974). When the animals were placed at 17 °C on a temperature gradient without food, young adults (day 1 of adulthood, D1) migrated up the temperature gradient toward 20 °C (Fig. 1B). On the other hand, aged animals (day 5 of adulthood, D5) remained around the spotted area and did not reach the area near T_cult_ (Fig. 1B), as previously reported (Huang et al., 2020). The ability to perform the thermotaxis behavior was quantified using the performance index, indicating the fraction of animals around T_cult_ (Fig. 1A). The performance index declined from D1 to D5 (Fig. 1C). To further accelerate aging (Fig. S1) (Klass, 1977), we cultivated animals at 23 °C and placed them on a temperature gradient centered at 20 °C. In this condition, animals gradually lost the ability to move toward T_cult_ during aging (Fig. S2), and the performance index declined from ∼0.75 at D1 to ∼0.25 at D5 (Fig. 1D). This age-dependent thermotaxis decline was not specific to OP50-fed animals because animals fed another *E. coli* strain, HT115, also showed a similar decline (Fig. S3).

**Figure 1.**
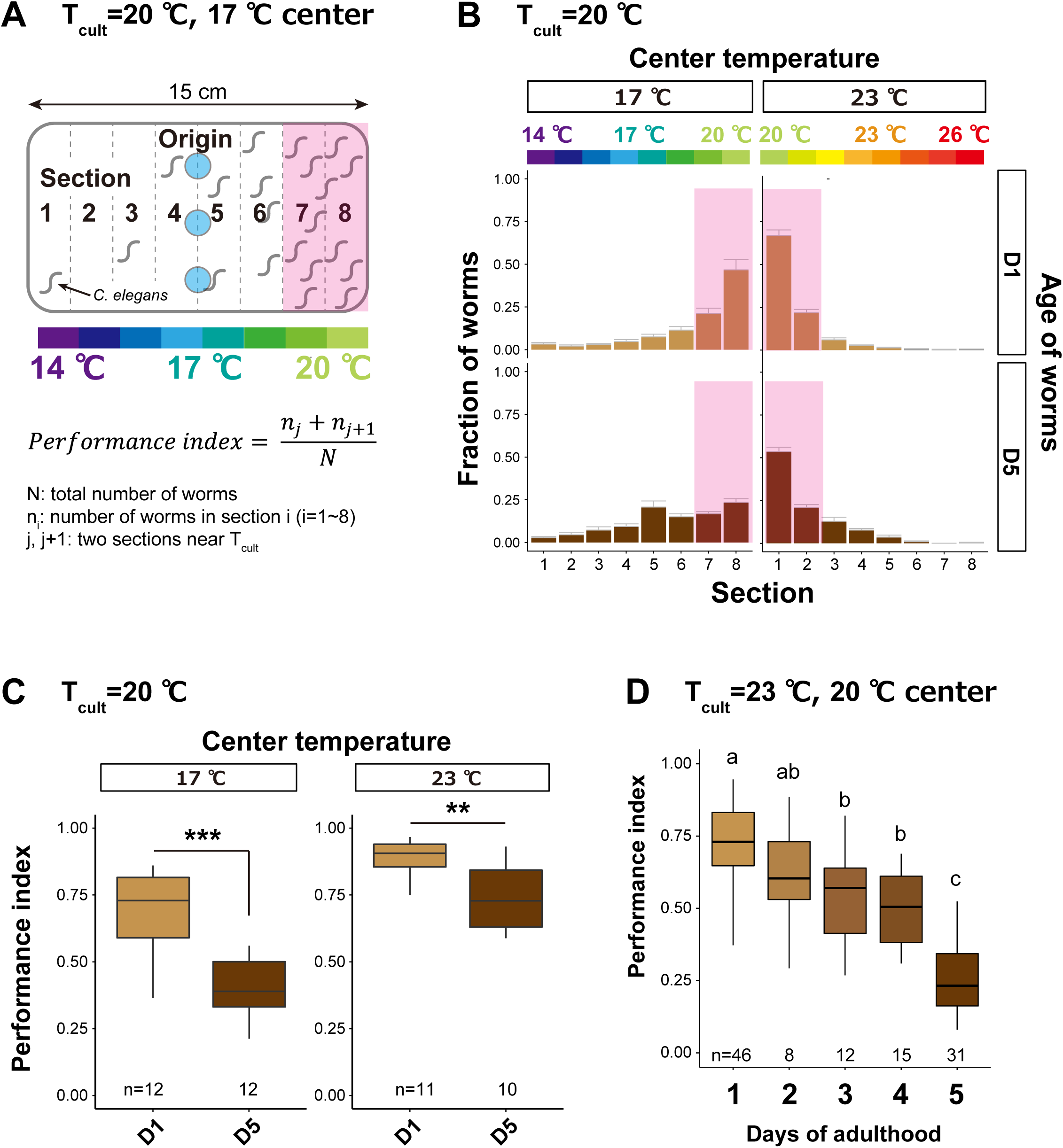
Thermotaxis performance declines with age. (A) Schematic of thermotaxis assay. Animals were placed at light blue circles on a thermal gradient without food. The pink rectangle indicates the sections around the T_cult_. After one hour, the number of animals in each section was counted to calculate the thermotaxis performance index using the indicated formula. (B and C) Age-dependent changes in thermotaxis behavior. D1 and D5 animals were cultivated with *E. coli* at 20 °C and placed at the center of a 14-20 or 20-26 °C gradient. (B) Distributions of animals (pink rectangle: the sections around the T_cult_) on the thermotaxis plates. (C) Box plots of thermotaxis performance indices. The number of experiments is shown. Statistics: Student’s t-test compared to D1 adults. **p<0.01, ***p<0.001. (D) Box plots summarizing thermotaxis performance indices of animals at different ages. Animals were cultivated with *E. coli* at 23 °C and placed at the center of a 17-23 °C gradient. The number of experiments is shown. Statistics: The mean indices marked with distinct alphabets are significantly different (p < 0.05) according to One-way ANOVA followed by Tukey– Kramer test.

The low thermotaxis performance of *E. coli*-fed aged animals appeared to be independent of defects in motility or temperature sensation because D5 animals cultivated at 20 °C could migrate down the thermal gradient relatively normally when the origin was at 23 °C (Figs. 1B and 1C). Consistent with this notion, we did not observe the loss of AFD or AIY neurons at D5 (Fig. S4). Moreover, aged animals could sense food normally based on the basal slowing response in the presence of food (Figs. S5A and S5B) (Sawin, Ranganathan, & Horvitz, 2000). The food sensation of aged animals is also reported to be normal, based on the attraction to *E. coli* (Cornils et al., 2016). In contrast to the basal slowing response, we note that aged animals did not show a significantly enhanced slowing response in the starved condition (Fig. S5B), implying that aged animals might not sense starvation normally.

To address if the defects can be observed in another associative learning behavior, we tested the salt avoidance behavior using two settings of assays (Saeki, Yamamoto, & Iino, 2001; Wicks, de Vries, van Luenen, & Plasterk, 2000) (Figs. S6A and S6B). As previously reported, naïve D1 animals were attracted by NaCl, while they avoided NaCl when cultivated with NaCl in the absence of food in both settings (Saeki et al., 2001; Wicks et al., 2000) (Figs. S6C and S6D). In contrast to the thermotaxis behavior, D5 animals showed normal salt-avoidance behaviors (Figs. S6C and S6D).

We thus concluded that age-dependent thermotaxis changes reflected the decline of specific associative learning behavior and determined to use it to examine dietary effects.

### Specific LAB prevent the age-dependent thermotaxis decline

To address if diets affect the age-dependent decline in thermotaxis, we fed animals with different lactic acid bacteria (LAB) species instead of the regular *E. coli*. We selected 35 LAB, consisting of 17 *Lactobacilli (Lb.)*, two *Pediococci (P.)*, two *Lactococci (Lc.)*, two *Streptococci (S.)*, five *Leuconostoc (Ls.)*, and seven *Bifidobacteria (B.)* (Table S1). To avoid developmental effects by feeding with LAB, we cultivated animals with *E. coli* until D1 before switching to LAB (Fig. 2A). Animals were cultivated at 23 °C and spotted at the center of the 17-23 °C gradient (Fig. 2A). Five LAB did not support the survival of animals during aging (Fig. 2B, NA). While eight LAB did not affect the thermotaxis performance indices of the aged animals compared to *E. coli*, 22 LAB significantly increased them (Fig. 2B). Among those, *P. pentosaceus*, *Lb. reuteri*, *Lb. rhamnosus*, and *Lb. plantarum* gave the highest performance indices (Fig. 2B).

**Figure 2.**
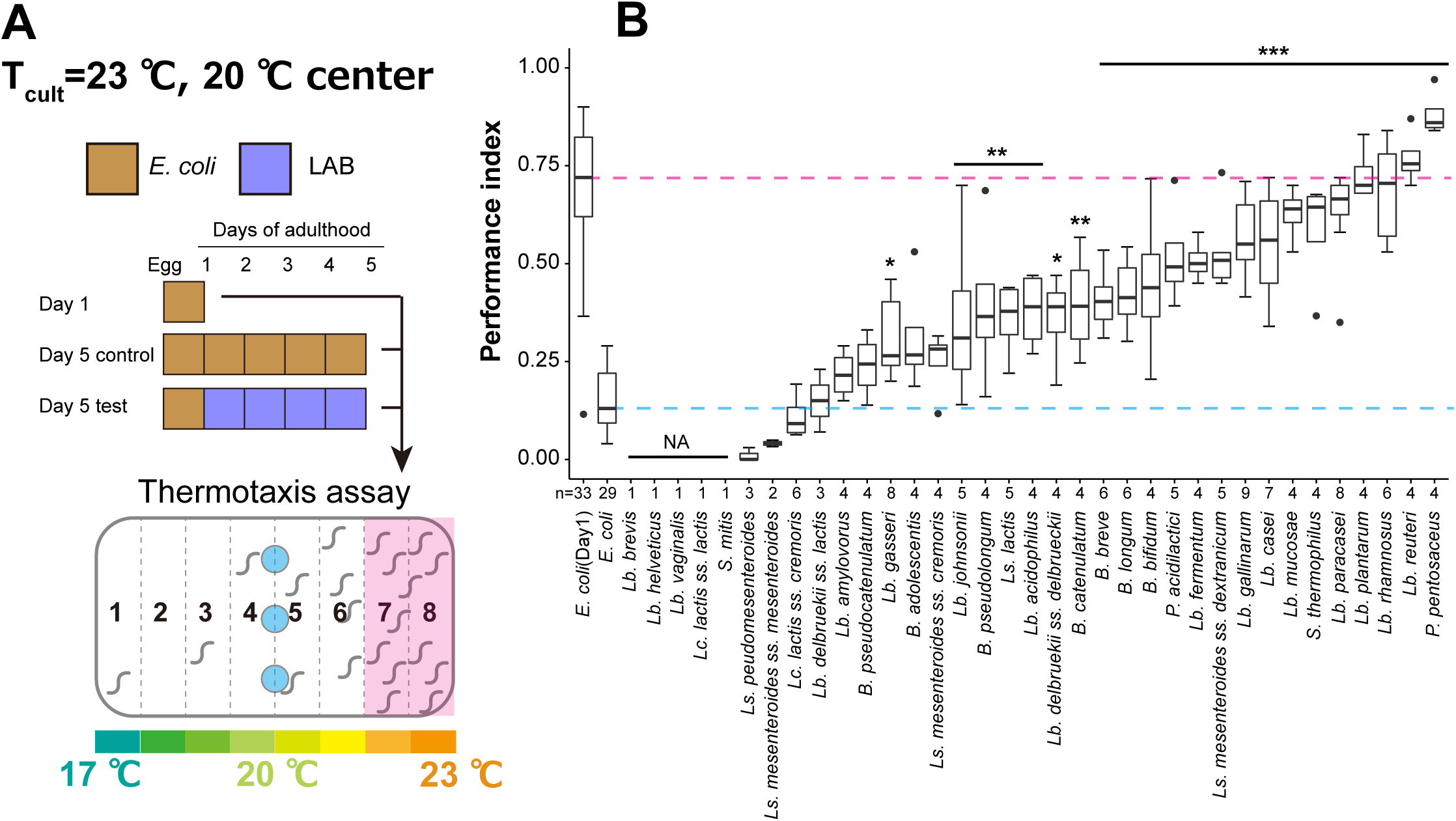
LAB screen for thermotaxis in aged animals. (A) Schematic of the screening procedure. Animals were cultivated at 23 °C with *E. coli* until D1 and transferred to *E. coli* or LAB plates every day until D5. At D5, animals were subjected to thermotaxis assays with a thermal gradient of 17-23 °C. (B) Box plots comparing thermotaxis performance indices of D5 animals fed LAB to those of D1 (pink dashed line) and D5 animals (light blue dashed line) fed *E. coli*. “Not applicable” (NA) indicates that animals fed those LAB were not subjected to the assay because they were sick or dead. Abbreviations: *B, Bifidobacterium; Lb, Lactobacillus; Lc: Lactococcus; Ls, Leuconostoc; P, Pediococcus; S, Streptococcus*. The number of experiments is shown. Statistics: One-way ANOVA followed by Dunnett’s multiple comparison test compared to D5 adults fed *E. coli*, ***p<0.001; **p<0.01; *p<0.05.

We first ruled out the possibility that aged animals fed the LAB were constitutively thermophilic, irrespective of the T_cult_. Thermophilicity is reported for mutants of genes such as *pkc-1*/*ttx-4* encoding protein kinase C (Okochi, Kimura, Ohta, & Mori, 2005) and *tax-6* encoding calcineurin A subunit (Kuhara, Inada, Katsura, & Mori, 2002). To distinguish between associative learning and thermophilicity, we shifted the thermal gradient of assay plates from 17-23 °C to 20-26 °C for animals cultivated at 23 °C (Fig. S7A). D5 *tax-6* mutants migrated toward a higher temperature than T_cult_ (Fig. S7B). On the other hand, LAB-fed D5 animals stayed around the T_cult_ (Fig. S7B). We calculated the thermotaxis index, instead of the performance index, to quantify animals’ thermal preference (Ito, Inada, & Mori, 2006) (Figs. S7A). Unlike thermophilic *tax-6* mutants, LAB-fed D5 animals showed thermotaxis indices comparable to the D1 wild type (Fig. S7C), suggesting that LAB-fed D5 animals were not constitutively thermophilic.

We next addressed if LAB-fed D5 animals can remember a new temperature by shifting the T_cult_ from 23 °C to 17 °C one day before the thermotaxis assay. D1 animals could learn the new temperature and shift their thermal preference from 23 °C (Fig. 3A) to the new T_cult_, 17 °C (Fig. 3B). The thermotaxis index was also shifted accordingly (Fig. 3C). LAB-fed D5 animals showed behavioral plasticity like D1 animals (Figs. 3B and 3C). This result suggests that LAB-fed aged animals retained an ability to learn a new T_cult_.

**Figure 3.**
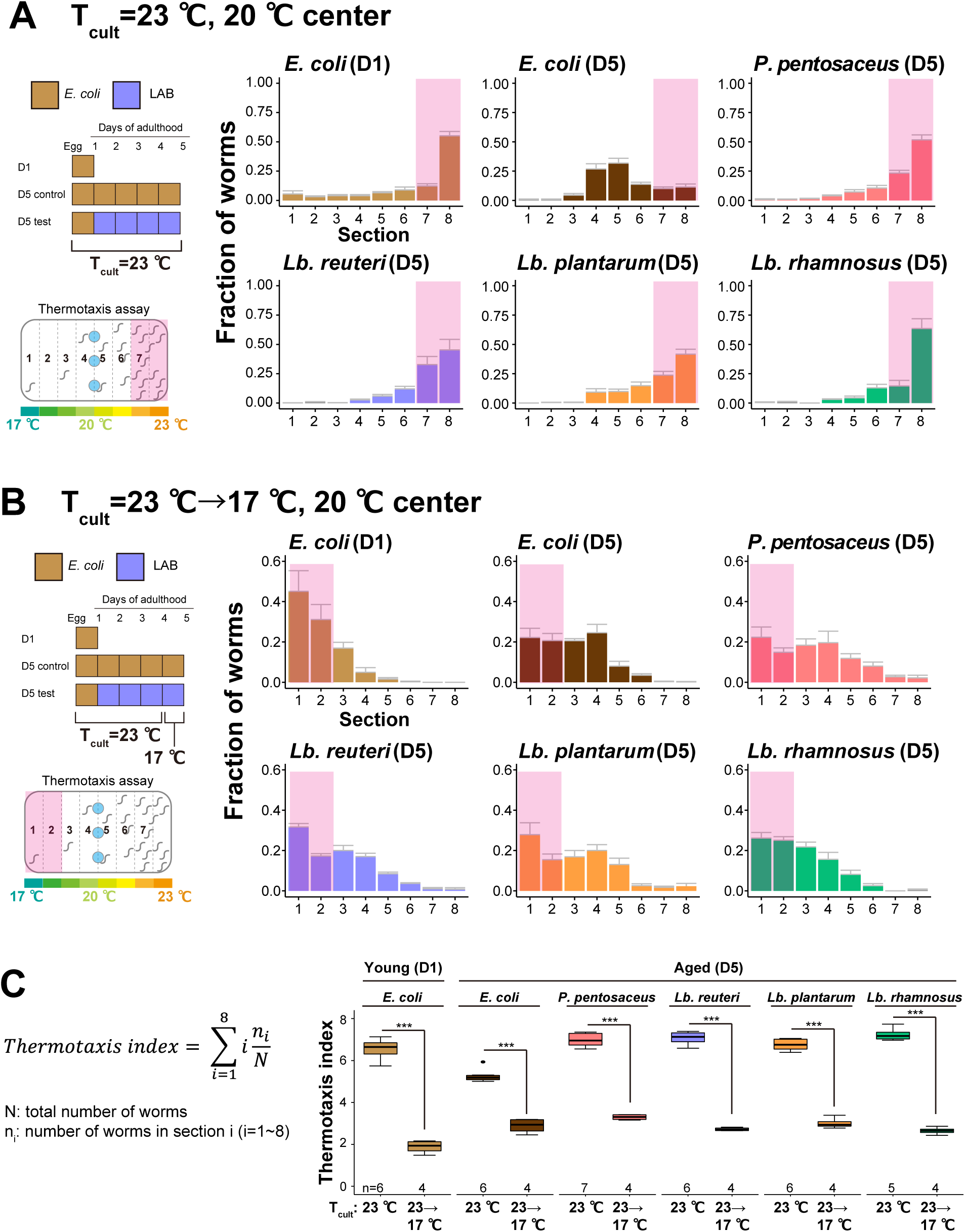
LAB-fed aged animals learn a new T_cult_. (A and B) The distribution of animals on thermotaxis plates. Pink rectangles indicate the sections around the T_cult_. (A) D1 or D5 animals fed indicated bacteria were cultivated at 23 °C and placed at the center of a 17-23 °C gradient. (B) Temperature shift assay. T_cult_ was shifted from 23 °C to 17 °C one day before the assay. Animals were placed at the center of the 17-23 °C gradient. (C) Box plots summarizing thermotaxis indices corresponding to (A) and (B). Thermotaxis indices were calculated to examine the mean distribution of animals on thermotaxis plates using the indicated formula. The number of experiments is shown. Statistics: Student’s t-test for comparison between T_cult_=23 °C and T_cult_=23 °C → 17 °C, ***p<0.001.

### Different LAB show various effects on lifespan and locomotion

Some LAB extend the lifespan of *C. elegans* (Komura et al., 2013; Nakagawa et al., 2016; Wang et al., 2020; Y. Zhao et al., 2013). Therefore, better thermotaxis performance of LAB-fed aged animals might result from a systemic effect of prolonged organismal lifespan. To address this possibility, we measured the lifespan of animals fed the four LAB: *P. pentosaceus*, *Lb. reuteri*, *Lb. rhamnosus*, and *Lb. plantarum.* To avoid the growth of *E. coli* on LAB plates after transferring animals, we used peptone-free NGM plates (T. Ikeda, Yasui, Hoshino, Arikawa, & Nishikawa, 2007; Lee, Kwon, & Lim, 2015). The lack of peptone in the culture plates did not affect the dietary effects on the thermotaxis of aged animals (Fig. S8). LAB had various effects on the lifespan of animals: *P. pentosaceus* prolonged the lifespan; *Lb. reuteri* did not affect the lifespan; *Lb. rhamnosus* and *Lb. plantarum* shortened the lifespan (Fig. 4A).

**Figure 4.**
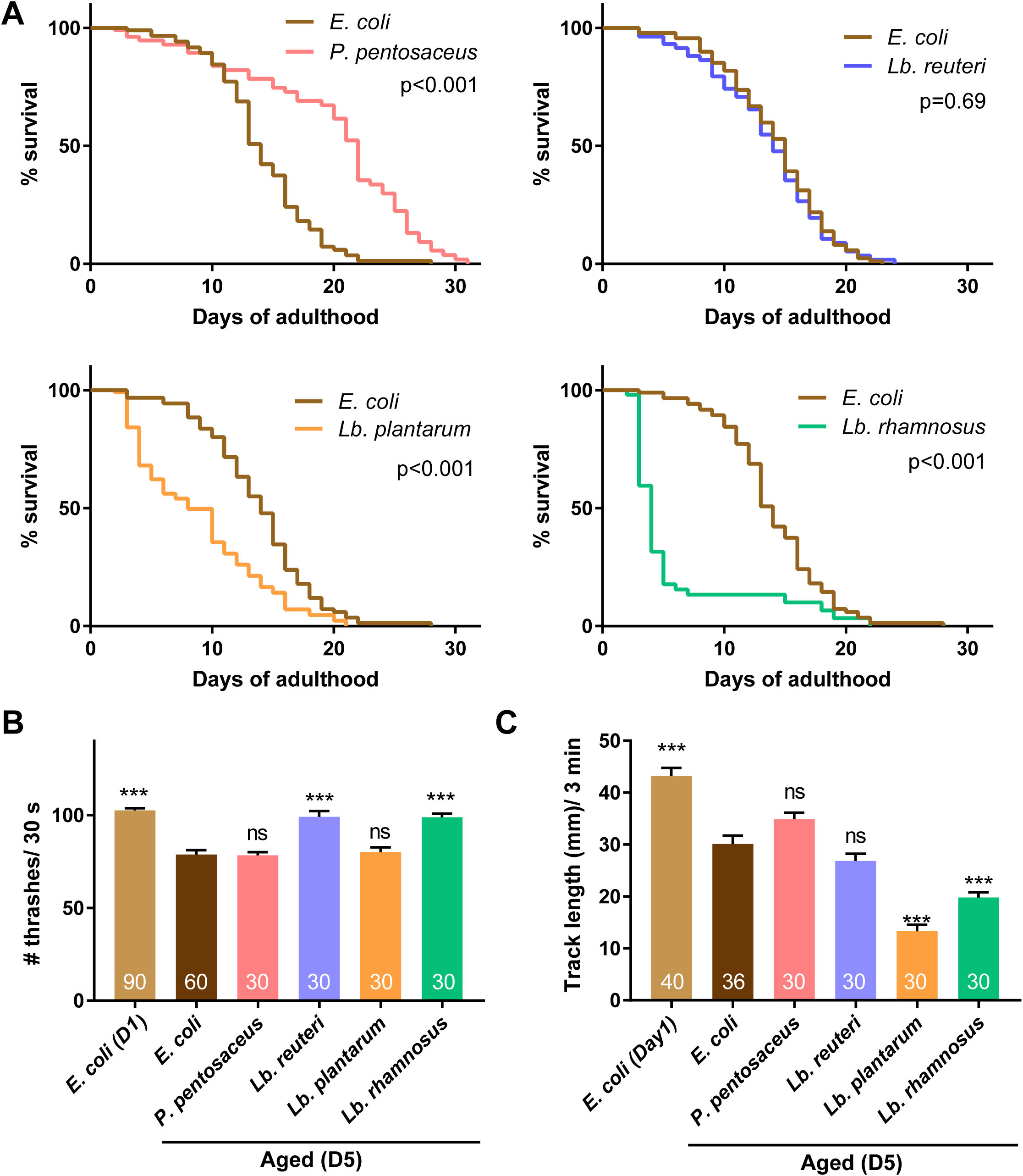
LAB shows various effects on lifespan and locomotion. Animals were fed *E. coli* until D1 and indicated bacteria after D1. (A) Survival curves of animals fed indicated LAB are shown with control animals fed *E. coli*. NGM plates without peptone were used to avoid the undesired growth of *E. coli* on LAB plates. N=4 experiments with 25 animals/experiment (100 animals in total). Statistics: Log-rank test. p values are shown. (B and C) The number of thrashes in liquid (B) and distance of migration in three minutes on plates with *E. coli* (C) were measured to examine the locomotion of aged animals. The number of animals is shown in bars. Error bars: S.E.M. Statistics: One-way ANOVA followed by Dunnett’s multiple comparison test compared to D5 fed *E. coli*, p***<0.001; ns, p>0.05.

We next examined the effect of LAB on the locomotion of aged animals using two assays: thrashing assay (Miller et al., 1996) and motility assay. As previously reported (Glenn et al., 2004; Hahm et al., 2015; Mulcahy et al., 2013), aged animals fed *E. coli* showed slight locomotion defects in both assays (Figs. 4B and 4C). In the thrashing assay, *Lb. reuteri*- and *Lb. rhamnosus*-fed aged animals showed better locomotion than *E. coli*-fed aged animals, while *P. pentosaseus* and *Lb. plantarum* did not have effects (Fig. 4B). In the motility assay, we measured the distance animals migrate on a plate. *Lb. plantarum-* and *Lb. rhamnosus*-fed aged animals showed reduced locomotion than *E. coli*-fed aged animals, while *P. pentosaseus* and *Lb. reuteri* did not have effects (Fig. 4C). Thus, the four LAB selected based on thermotaxis had different effects on the lifespan and locomotion, implying that the dietary effect on thermotaxis is independent of lifespan and motility.

### Bacteria affect the age-dependent thermotaxis decline as nutrition

How do different bacteria affect the thermotaxis of aged animals? *C. elegans* shows different preferences in the bacterial diet (Shtonda & Avery, 2006). Our LAB screen used different diets during the temperature-food association before the thermotaxis assays (Fig. 2A). It raises the possibility that the different strengths of the association during learning caused the difference in thermotaxis of aged animals. To address this issue, we switched the foods one day before the thermotaxis assay (Fig. 5A). We used *Lb. reuteri* because it did not affect the lifespan (Fig. 4A). Aged animals whose diet was switched from *Lb. reuteri* to *E. coli* showed the high thermotaxis performance, while aged animals with the opposite condition did not (Fig. 5A). This result suggests that the dietary effects of thermotaxis on aged animals do not reflect the strength of association.

**Figure 5.**
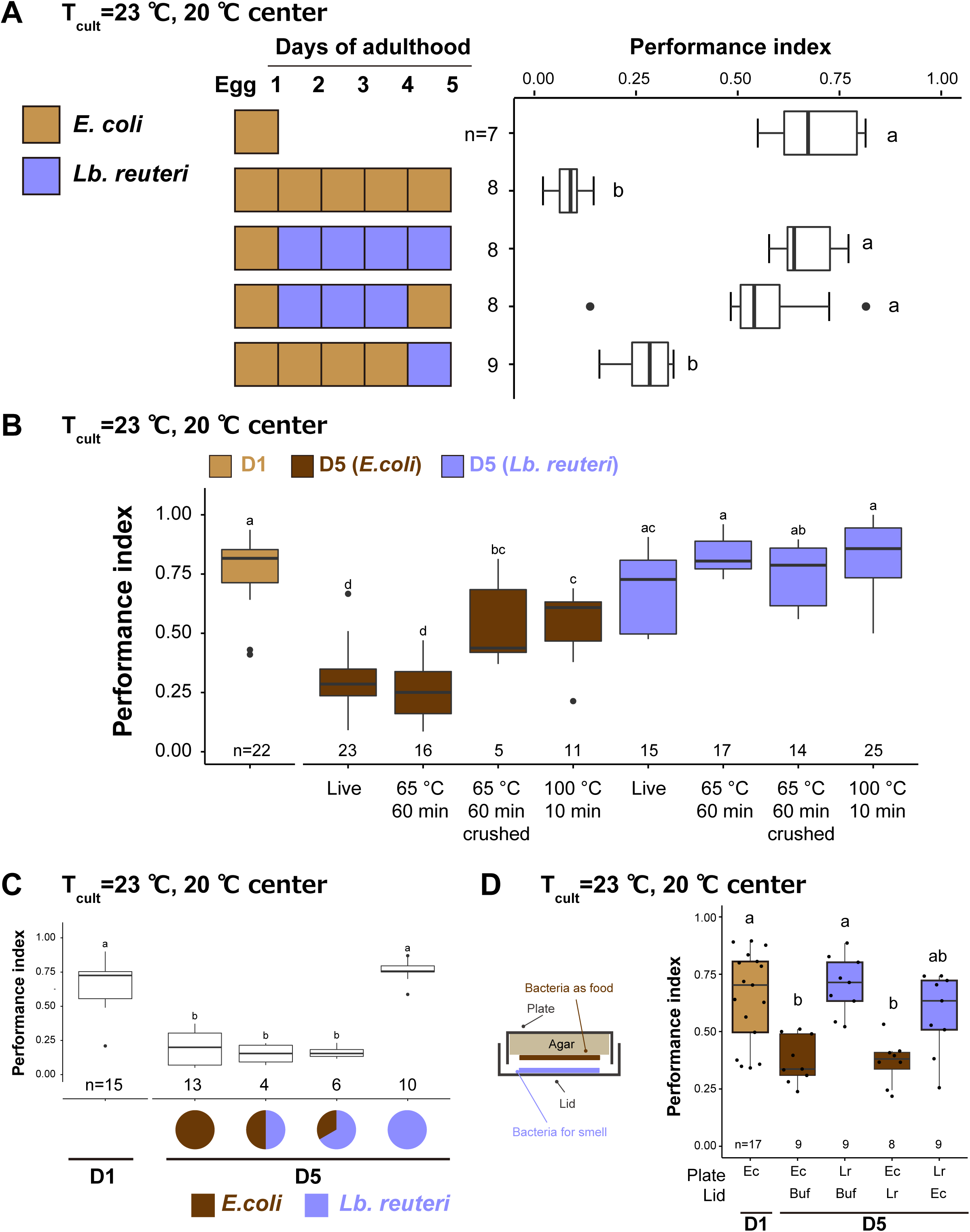
Bacteria affect thermotaxis of aged animals as nutrition. Box plots show thermotaxis performance indices of animals fed indicated bacteria and cultivated at 23 °C. Aged animals were transferred every day to new plates from D1. The number of experiments is shown. Statistics: The mean indices marked with distinct alphabets are significantly different (p < 0.05) according to One-way ANOVA followed by Tukey–Kramer test. (A) The short-term effects of diet. The diet was switched one day before the thermotaxis assay, as indicated in the schematic. (B) The effect of heat treatment and crushing of bacteria. The bacteria were killed by incubating at 65 °C for 1 hour or 100 °C for 10 min. After heat treatment, bacteria were crushed using a bead-based homogenizer. (C) The mixture of bacteria. Live *E. coli* and *Lb. reuteri* were mixed at a 1:1 or 1:2 ratio with the final concentration of 0.1 g/ml and used as a diet. (D) The effect of bacterial odor. Animals were exposed to the bacterial odor by putting the bacterial solution on the lid and cultivated, as shown in the schematic. Ec: *E. coli*, Lr: *Lb. reuteri*.

To examine if LAB affects animals as live bacteria or serves as nutrition, we examined the effect of bacteria heat-killed at 65 °C for one hour or at 100 °C for ten minutes (see Materials and Methods). Like aged animals fed live bacteria, ones fed 65 °C-treated *E. coli* and *Lb. reuteri* showed low and high performance indices in thermotaxis, respectively (Fig. 5B). Thus, the effect of both bacteria on thermotaxis is independent of the condition of the bacteria being alive. On the other hand, animals fed 100 °C-treated *E. coli* showed higher performance than those fed live *E. coli* (Fig. 5B). This result suggests that a component in *E. coli* resistant to 65 °C but sensitive to 100 °C facilitates thermotaxis decline during aging. Moreover, animals fed crushed *E. coli* also showed higher thermotaxis performance, implying that the said component might be diffused after crushing (Fig. 5B).

Next, we examined which bacteria, *E. coli* or *Lb. reuteri* has dominant effects on the thermotaxis of aged animals. We mixed *E. coli* and *Lb. reuteri* and fed animals from D1. Both *E. coli* and *Lb. reuteri* were ingested by *C. elegans* even when mixed, based on FITC labeling of bacteria (Fig. S9). In this condition, *E. coli* had a dominant effect even when *Lb. reuteri* was mixed with twice as much as *E. coli* (Fig. 5C). The dominant effect of *E. coli* seemed to require ingestion of bacteria because the exposure to the *E. coli* odor did not affect the thermotaxis ability of *Lb. reuteri*-fed aged animals (Fig. 5D).

Collectively, we concluded that ingestion of bacterial nutrition during aging affects the thermotaxis of aged animals.

### The effects of LAB are associated with the phylogenetic tree

We explored the different features of bacteria which might affect *C. elegans* physiology. Gram-staining showed that *E. coli* was Gram-negative, while LAB were Gram-positive as expected (Fig. S10). The morphologies and sizes of the four select LAB were different (Fig. S10); these physical properties of LAB may not explain the high performance indices of aged animals.

To examine the features of the genus of LAB, we analyzed a phylogenetic tree of 35 LAB strains with the heatmap of the associated thermotaxis performance indices (Fig. 6). This analysis revealed that LAB associated with high performance indices was significantly enriched in a specific clade, henceforth referred to as Clade A (Fig. 6). Clade A, including *Lactobacilli* and *Pediococci,* was enriched in obligatory and facultatively heterofermentative species except for *P. pentosaceus*, which was obligatory homofermentative (Fig. 6). On the other hand, the *Lactobacilli*, associated with relatively low thermotaxis indices (Clade B), are all homofermentative (Fig. 6). These results suggest that the nutritional feature of LAB shared among the clades of *Lactobacilli* might affect *C. elegans*.

**Figure 6.**
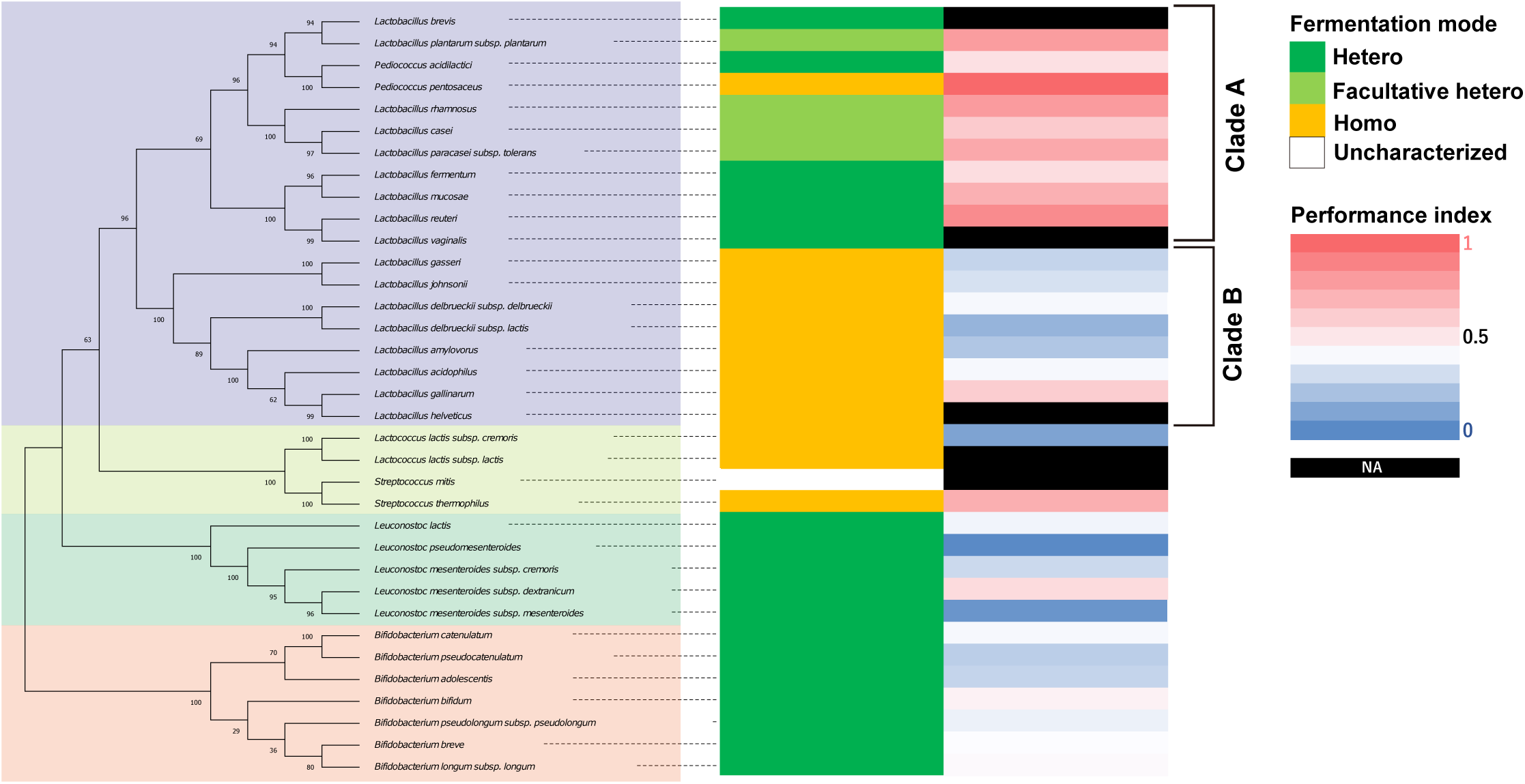
L*a*ctobacilli in a clade are associated with high thermotaxis performance of aged animals. Phylogenetic tree of LAB based on 16S rRNA is shown with fermentation mode, and heatmap of performance indices of aged animals fed indicated LAB from D1. Bootstrap values are indicated at each node on the phylogenetic tree. Fermentation modes were categorized based on previous studies (see Table S1). The same data as Figure 2B were used for thermotaxis performance indices. NA in the performance indices heatmap indicates that animals fed those LAB were not subjected to the thermotaxis assay because they were sick or dead. Fermentation mode indicates obligatory hetero- (green), facultatively hetero- (light green), and obligatory homofermentative (orange) LAB.

### Neuronal *daf-16* is involved in the thermotaxis performance of *Lb. reuteri*-fed aged animals

Since the dietary effect on thermotaxis required more than one day to be manifested, bacterial diets seem to alter the internal state of *C. elegans*. We addressed the molecular mechanism of how different diets affect *C. elegans*. LAB can induce dietary restriction, which leads to a prolonged lifespan (Y. Zhao et al., 2013). However, three out of four select LAB did not increase lifespan (Fig. 4A). *pha-4*, an ortholog of the human FOXA2 transcription factor, is required for dietary restriction-induced longevity, and its expression is increased by dietary restriction (Panowski, Wolff, Aguilaniu, Durieux, & Dillin, 2007). In our condition, *pha-4* expression decreased in LAB-fed aged animals compared to *E. coli*-fed aged animals, suggesting that LAB-fed animals were not under dietary restriction (Fig. S11A). To directly test the effect of dietary restriction on the thermotaxis, we used *eat-2* mutants, which exhibit dietary restriction by defective pharyngeal pumping (Lakowski & Hekimi, 1998). *eat-2* mutants did not increase the performance index of *E. coli*-fed aged animals, although it may be due to the involvement of *eat-2* in thermotaxis as shown in D1 animals (Fig. S9B). These results suggest that the high thermotaxis performance of LAB-fed aged animals is not due to dietary restriction.

To find the gene involved in the dietary effect on aged animals, we tested mutants of three genes: *nkat-1* and *kmo-1* genes that encode enzymes in the kynurenic acid synthesizing pathway are known to be involved in butanone-associated memory in aged animals (Vohra et al., 2018); *daf-16* that is an ortholog of mammalian FOXO transcription factor involved in longevity (Kenyon et al., 1993) and LAB-dependent lifespan extension (Grompone et al., 2012; Lee et al., 2015; Sugawara & Sakamoto, 2018). Aged *nkat-1* and *kmo-1* mutants maintained thermotaxis ability like wild type when fed *Lb. reuteri*. On the other hand, aged *daf-16* mutants showed significantly less ability to perform thermotaxis than their D1 counterpart (Fig. 7A); aged *daf-16* mutants fed *Lb. reuteri* distributed around a temperature slightly lower than the T_cult_ (Fig. 7B). This decreased thermotaxis ability in aged *daf-16* mutants fed *Lb. reuteri* was not due to shortened lifespan because *daf-16* mutants had comparable lifespan to wild-type animals when fed *Lb. reuteri* (Fig. 7C). DAF-16 is known to be activated in *daf-2* mutants. Therefore, we speculated that *daf-2* mutants might show high thermotaxis performance even in *E. coli*-fed aged animals. However, *daf-2* was required for nomal thermotaxis both young and aged animals (Fig. 7D).

**Figure 7.**
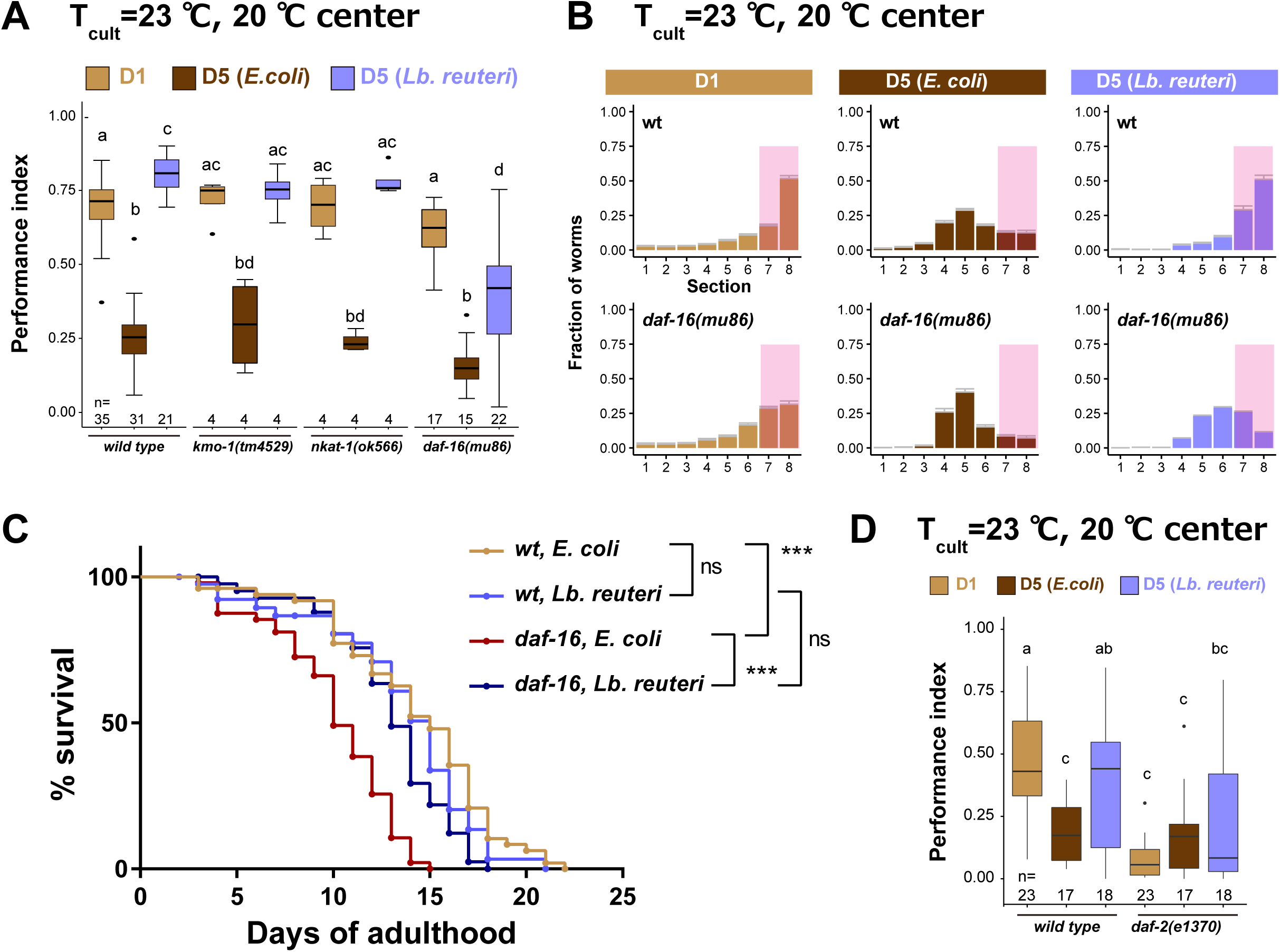
d*a*f*-16* is involved in the effect of *Lb. reuteri* on thermotaxis in aged animals. (A) Box plots summarizing thermotaxis performance indices of animals with indicated genotypes. Animals were cultivated at 23 °C with *E. coli* or *Lb. reuteri* from D1. The number of experiments is shown. Statistics: One-way ANOVA followed by Dunnett’s multiple comparison test compared to the D1 control for each condition, p***<0.001; ns, p>0.05. (B) Distribution of animals of indicated conditions on thermotaxis plates. Pink rectangles indicate the sections around the T_cult_. (C) Survival curves of animals with indicated genotypes fed *E. coli* or *Lb. reuteri* from D1 and cultivated at 23 °C. NGM plates without peptone were used. N=4 experiments with 25 animals/experiment (100 animals in total). Statistics: Log-rank test, p***<0.001; ns, p>0.05. (D) Box plots summarizing thermotaxis performance indices of animals with indicated genotypes. Animals were cultivated at 15 °C for 96 hours to avoid dauer formation of *daf-2(e1370)* and then incubated at 23 °C until D1 or D5 with *E. coli* or *Lb. reuteri*. The number of experiments is shown. Statistics: The mean indices marked with distinct alphabets are significantly different (p < 0.05) according to One-way ANOVA followed by Tukey–Kramer test.

*daf-16* possesses several isoforms with different expression patterns and functions (Fig. 8A) (Kwon, Narasimhan, Yen, & Tissenbaum, 2010). The b isoform is a neuronal isoform involved in AIY development (Christensen, de la Torre-Ubieta, Bonni, & Colon-Ramos, 2011). The other isoforms are more broadly expressed in most tissues, including neurons. We addressed which isoform is necessary for the effect on thermotaxis of aged animals. *daf-16(mg54)* has a single nucleotide polymorphism (SNP) which introduces an amber stop codon mutation and affects the exons of all *daf-16* isoforms except the b isoform (Fig. 8A) (Ogg et al., 1997). To knock out only the b isoform, we generated *daf-16(knj36)* which introduced 7-bp deletion in the first exon of the b isoform, located in the intron of the other isoforms (Fig. 8A). Both *daf-16(mg54)* and *daf-16(knj36)* did not affect the dietary effects on the thermotaxis of aged animals (Fig. 8B). This result implies that the b isoform and other isoforms complement each other and that having either one is sufficient to give the dietary effect. Consistently, the expression of *daf-16b* under the control of its own promoter rescued the deletion mutants, *daf-16(mu86),* confirming the sufficiency of the *daf-16 b* isoform.

**Figure 8.**
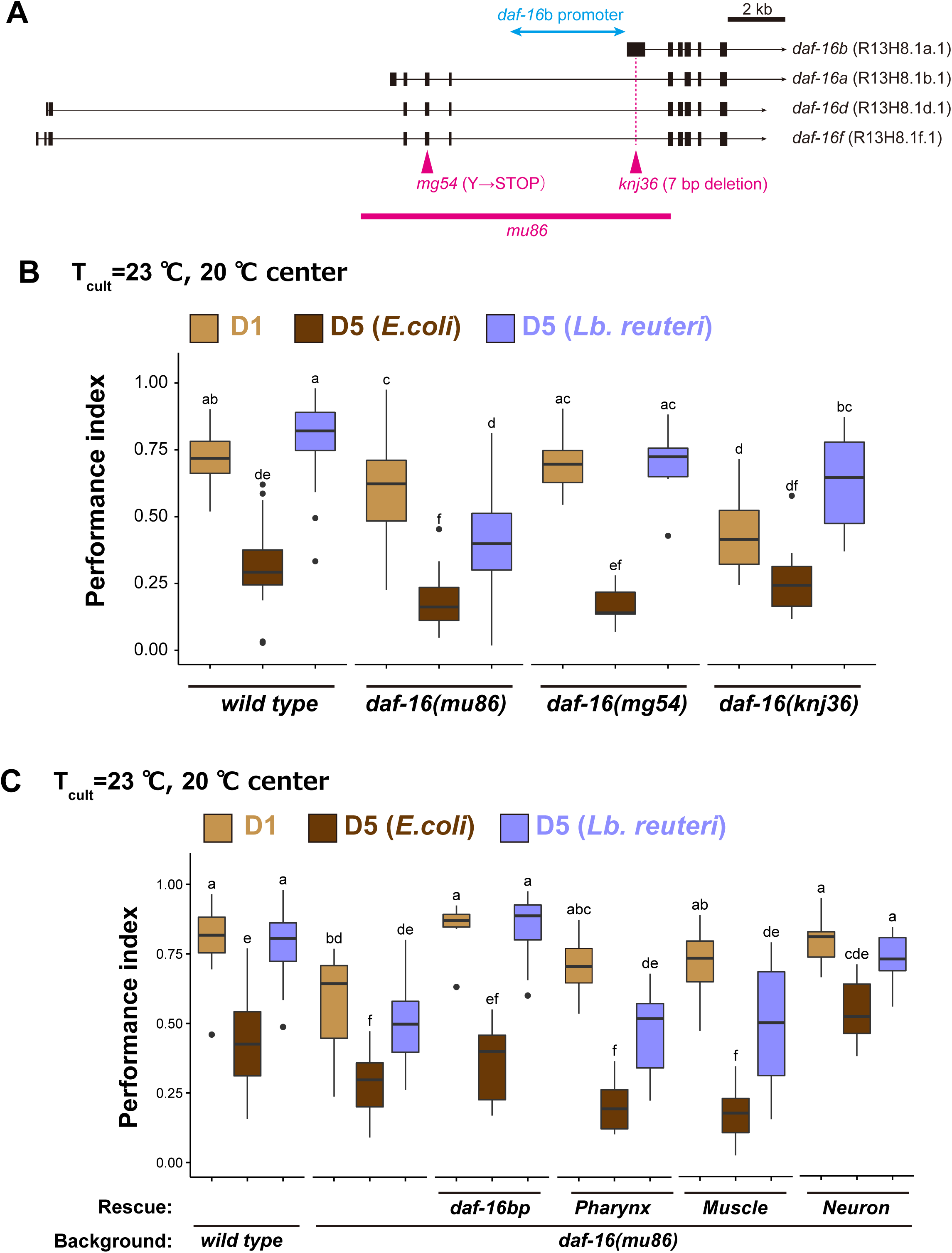
d*a*f*-16* b isoform functions in neurons to maintain high thermotaxis ability in aged animals fed *Lb. reuteri*. (A) Schematic of *daf-16* locus with representative isoforms based on WormBase (https://wormbase.org). Black boxes and black lines indicate exons and introns, respectively. Arrows indicate 3’UTR. Alleles used in this study are shown in magenta. The promoter of the *daf-16* b isoform (4.9 kbp) is indicated in light blue. (B and C) Box plots summarizing thermotaxis performance indices of animals with indicated genotypes. Animals were cultivated at 23 °C with *E. coli* or *Lb. reuteri* from D1. The number of experiments is shown. Statistics: The mean indices marked with distinct alphabets are significantly different (p < 0.05) according to One-way ANOVA followed by Tukey–Kramer test. (B) Analysis of different alleles of *daf-16* indicated in (A). (C) Tissue-specific rescue. The single copy-insertions of *daf-16*b fragment with introns under tissue-specific promoters were used to examine if it rescues *daf-16(mu86)*: *myo-2*p, pharynx; *myo-3*p, body-wall muscle; *rgef-1*p, pan-neuron.

We addressed in which tissue *daf-16* functions. Given that the expression of either b isoform or non-b isoforms was sufficient to provide the dietary effects (Fig. 8B), *daf-16* might function in a tissue where both b and non-b isoforms are expressed. We thus focused on pharyngeal muscles, body wall muscles, and neurons (Nagashima, Iino, & Tomioka, 2019). Expression of *daf-16b* in pharyngeal muscles or body wall muscles did not rescue *daf-16(mu86)* (Fig. 8C). On the other hand, the *daf-16 b* isoform expressed in all neurons rescued the low performance of *Lb. reuteri*-fed aged *daf-16(mu86)* mutants (Fig. 8C). This result suggests that *daf-16* functioning in neurons is involved in the high thermotaxis performance of *Lb. reuteri*-fed aged animals.

### Diet and age affect DAF-16 target genes

To comprehensively understand the dietary effect on aging, we carried out RNA sequencing of eight samples: D1, D5 fed *E. coli*, D5 fed homo-fermentative LAB, which gave low thermotaxis index (*Lb. gasseri* and *Lb. delbrueckii*), and D5 fed heterofermentative LAB, which gave high thermotaxis index (*P. pentosaceus*, *Lb. reuteri*, *Lb. rhamnosus*, and *Lb. plantarum*) (Fig. 9A). Principal component analysis of transcriptome data revealed that the D5 fed LAB clustered together separately from D1 and D5 fed *E. coli*, irrespective of the fermentation mode of LAB (Fig. 9B). The first principal component (PC1) explained 76% of the entire variance and appeared to represent the difference in age irrespective of diets (Fig. 9B, PC1). To characterize what genes contributed to PC1, we selected the top 5% genes positively or negatively correlating to PC1 (666 genes, each, among 13331 genes in total, Fig. S12A) and performed Gene Ontology (GO) analysis using Metascape (Y. Zhou et al., 2019). The genes correlating to PC1 were enriched with the categories such as oogenesis (GO:0048477; p=1.0×10^-10^; Enrichment, 5.7) and structural constituent of cuticle (GO:0042302; p=1.0×10^-34^; Enrichment, 8.3) (Fig. S12C). The second principal component (PC2) explained 9% of the entire variance and appeared to represent the difference between *E. coli* and LAB (Fig. 9B). GO analysis showed that the genes correlating to PC2 were enriched with the categories such as glucuronosyltransferase activity (GO:0015020; p=1.0×10^-10^; Enrichment, 6.7) and biological process involved in interspecies interaction between organisms (GO:0044419; p=1.0×10^-34^; Enrichment, 5.3) (Figs. S12B and S12D). Among these categories, the neuropeptide signaling pathway (GO:0007218; p=5.1×10-4; Enrichment, 2.9) is particularly intriguing because it can work as signals from the intestine to neurons.

**Figure 9.**
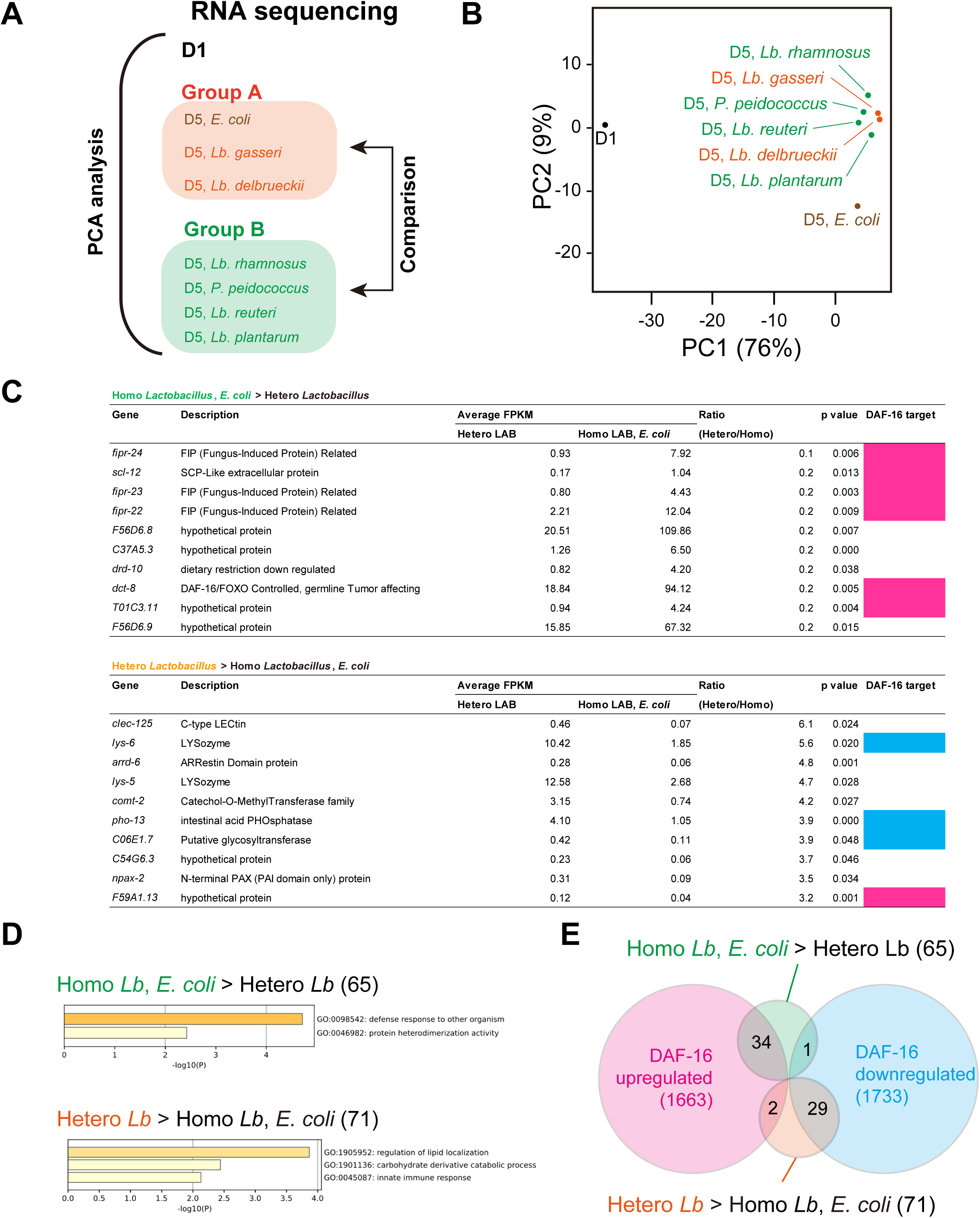
Transcriptome analysis on the effect of aging and diet. (A) Schematic of transcriptome analysis. We carried out RNA sequencing of indicated samples (eight in total). Principal component analysis was carried out using all samples. Differentially expressed genes were analyzed using D5 samples to examine the difference between Group A and Group B. (B) A scatter plots of the two variables projected on the first and second principal components. The percentage of variance explained by each principal component is indicated in brackets. (C) Differentially expressed genes (DEG) between Group A and Group B in (A). DEG were ranked by the ratio between Group A’s and B’s average expression, and the top 10 genes are shown. Magenta and light blue indicate genes that are upregulated and downregulated by DAF-16 (Tepper et al., 2013), respectively. (D) Gene ontology analysis of differentially expressed genes between Group A and B. x-axis indicates log_10_[p-value of enrichment]. The number of genes is indicated in brackets. (E) Venn diagram showing the overlap between differentially expressed genes of our samples and DAF-16 targets (Tepper et al., 2013). The number of genes is indicated in brackets.

The principal component analysis revealed the feature of gene expression of animals of different ages and diets. However, it did not explain the difference in thermotaxis ability in (1) D5 animals fed *E. coli* or homo fermentative LAB and (2) D5 animals fed hetero fermentative LAB. Therefore, we investigated differentially expressed genes between these two groups (Fig. 9A). We found 65 genes whose expression is >2 times higher in Group 1 (*E. coli* and Homofermentative LAB) than in Group 2 (Heterofermentative LAB). These genes included fungus-induced protein-related genes (*fipr-22*, *fipr-23*, and *fipr-24*) (Fig. 9C) and were enriched with the category such as regulation of lipid localization (GO:1905952; p=1.3×10^-4^; Enrichment, 29.0) (Fig. 9D). On the other hand, we found 71 genes whose expression is >2 times higher in Group 2 than in Group 1. These genes included lysozyme genes (*lys-5* and *lys-6*) (Fig. 9C) and were enriched with the category such as regulation of defense response to other organisms (GO:0098542; p=2.0×10^-5^; Enrichment, 6.6) (Fig. 9D).

Since the high thermotaxis ability of *Lb. reuteri*-fed aged animals were *daf-16*-dependent, we investigated the enrichment of *daf-16* targets in the whole body (1663 upregulated genes and 1733 downregulated genes (Tepper et al., 2013)). The genes highly expressed in aged animals fed *E. coli* or homofermentative LAB were significantly enriched with upregulated genes by DAF-16 (Normalized Enrichment Score (NES), -1.34; False Discovery Rate (FDR), 0.11; p<0.001) (Fig. 9E). On the other hand, the genes highly expressed in heterofermentative LAB were significantly enriched with downregulated genes by DAF-16 (NES, 1.53; FDR, 0.046; p<0.001) (Fig. 9E). This result implied that DAF-16 might be activated in aged animals fed *E. coli* or homofermentative LAB. Interestingly, this is the opposite of the prediction based on the *daf-16* dependency of the thermotaxis of *Lb. reuteri*-fed animals. Namely, *daf-16* is predicted to be activated in heterofermentative *Lb. reuteri*-fed aged animals. Therefore, *Lb. reuteri* might activate DAF-16 in a tissue and/or target-specific manner. Consistent with this notion, the general activation of DAF-16 using *daf-2* mutants did not improve the thermotaxis ability of aged animals (Fig. 7D).

## Discussion

Addressing the causal relationship between diets and their effects on animals’ physiology is challenging in human or mammalian models because microbiota in the gut and diet are complex. It is especially true in the context of aging because of their long lifespan. Using *C. elegans* as a model, we provide evidence of the dietary effect on the age-dependent behavioral decline discernible from the lifespan.

### Aging and diet show various effects on different behaviors and organismal lifespan

This study demonstrated that diets affect the age-dependent decline of thermotaxis behavior. We ruled out the possibility that the high performance of LAB-fed aged animals was due to thermophilicity, stronger association to LAB, better motility, dietary restriction, or longer lifespan. Thus, aging and diets likely affect the thermosensory circuit. The primary thermosensory neuron AFD (Mori & Ohshima, 1995) cell-autonomously stores temperature memory (Kobayashi et al., 2016). Although the Ca^2+^ response in AFD is reported to be defective in aged animals (Huang et al., 2020), the temperature sensation itself does not seem to be abolished in aged animals because they could migrate down the gradient (Figs. 1B and 1C). AFD thermosensory neurons synapse onto and regulate AIY interneurons by switching excitatory and inhibitory signals in a context-dependent manner (Mori & Ohshima, 1995; Nakano et al., 2020; White, Southgate, Thomson, & Brenner, 1986). AIY neurons are reported to be a cite of action of *age-1* PI3 kinase, which is upstream of *daf-16* in isothermal tracking behavior (H. Murakami et al., 2005). Therefore, aging and diet might affect AIY interneuorns.

The degree of age-dependent decline seems to depend on behaviors. *E. coli-*fed animals experienced an age-dependent decline in thermotaxis but not in salt avoidance behavior. Despite the similar effects of *P. pentosaceus, Lb. reuteri, Lb plantarum,* and *Lb. rhamnosus* on thermotaxis in aged animals, these LAB showed various effects on locomotion. In thermotaxis, aged animals showed more severe defects in migrating up the thermal gradient than migrating down (Figs. 1B and 1C). Thermotaxis is achieved by multiple steps: sensing temperature, recognizing food, associating food and temperature, memorizing T_cult_, and migrating toward T_cult_ (Aoki & Mori, 2015; Goodman & Sengupta, 2018; Kimata, Sasakura, Ohnishi, Nishio, & Mori, 2012). Thus, the different severities of thermotaxis decline between migration up and down the gradient in aged animals might be attributed to the different neural circuits responsible for those conditions, as previously reported (M. Ikeda et al., 2020).

Neuronal aging is discernible from an organismal lifespan. *nkat-1* mutants prevent age-dependent memory decline in associative learning between food and butanone without changing lifespan (Vohra et al., 2018). Similarly, we found that *Lb. reuteri* improved thermotaxis in aged animals without changing their lifespan. More strikingly, *Lb plantarum* and *Lb. rhamnosus* shortened the lifespan while they had beneficial effects on the thermotaxis of D5 adults. This different dietary condition will allow us to address the mechanism underlying phenotypic variation in aged animals independent from organismal lifespan and genetic perturbation.

### Diets affect age-dependent thermotaxis decline as nutrients

Previous reports elucidated how bacterial diet affects *C. elegans* as nutritional components, gut microbiota, and/or pathogen (Kumar et al., 2019; J. J. Zhou, Chun, & Liu, 2019) probably by changing *C. elegans* metabolites (Gao et al., 2017; Reinke, Hu, Sykes, & Lemire, 2010) and gene expression (MacNeil, Watson, Arda, Zhu, & Walhout, 2013).

Both live *E. coli* and LAB can colonize animals (Berg et al., 2016; Chelliah et al., 2018; Park et al., 2018; Portal-Celhay, Bradley, & Blaser, 2012). Live bacteria are necessary for some physiological roles; secreted enterobactin from live *E. coli* in the gut promotes *C. elegans* growth (Qi & Han, 2018); live, but not dead, LAB reduces the susceptibility to pathogenic bacteria *Pseudomonas aeruginosa*. On the other hand, live bacteria are unnecessary in different contexts; heat-killed *Lb. paracasei* and *Bifidobacterium longum* extend *C. elegans* lifespan (Sugawara & Sakamoto, 2018; Wang et al., 2020). In our thermotaxis assay on aged animals, *E. coli* and LAB killed by 65 °C treatment had similar effects to live bacteria. This result implies that, instead of the action of live bacteria, bacterial nutrition might be responsible for the effect on the thermotaxis of aged *C. elegans*. Feeding mixed bacteria suggests that *E. coli* has a dominant effect over *Lb. reuteri* to reduce the thermotaxis ability of aged animals. *E. coli* appeared to have components that are vulnerable at 100 °C and diffusible after crushing. Since the smell of bacteria did not reverse the effect of diet, the ingestion of bacterial nutrition causes the dietary effects. Metabolites in bacterial diet affect *C. elegans* physiology; some metabolites are beneficial, while others are toxic (J. J. Zhou et al., 2019). Coenzyme Q in *E. coli* shortens the lifespan of *C. elegans* (Larsen & Clarke, 2002). Bacterial nitric oxide and folate are also positive and negative regulators of *C. elegans* lifespan, respectively (Gusarov et al., 2013; Virk et al., 2012). Vitamin 12 in *Comamonas aquatica* accelerates development and reduces fertility without changing lifespan (Watson et al., 2014). Given that different metabolites are produced by different LAB (Tomita, Saito, Nakamura, Sekiyama, & Kikuchi, 2017), these metabolites might be responsible for the different effects on the thermotaxis of aged *C. elegans*.

Our results indicated that LAB associated with high performance indices of thermotaxis are associated with a clade enriched in heterofermentative *Lactobacilli* and *Pediococci* (Clade A in Figure 5A). Heterofermentative LAB produce not only lactic acid and ATP but also several other end products such as ethanol and CO_2_ from glucose. On the other hand, homofermentative LAB converts glucose into two molecules of lactic acid and ATP. Heterolactic fermentation itself does not explain the high performance index in thermotaxis of aged animals because heterofermentative *Leuconostoc* and *Bifidobacteria* species did not give the high performance indices. Metabolites other than lactic acid, ethanol, and CO_2_ also differ between hetero- and homofermentative *Lactobacilli* (Tomita et al., 2017). Metabolites enriched in heterofermentative *Lactobacilli* include a neurotransmitter GABA and tyramine, a substrate to synthesize neurotransmitter octopamine; metabolites enriched in homofermentative *Lactobacilli* include 4-hydroxyphenyllactic acid (HPLA) and acetoin. We note that Tomita *et al*. reported the metabolites in the media (Tomita et al., 2017) while we supply bacteria to animals after washing off the bacterial media. Nonetheless, metabolites enriched in different *Lactobacilli* might affect the age-dependent thermotaxis decline.

### Bacterial diets modulate genetic pathways in *C. elegans* neurons

LAB can extend the lifespan of *C. elegans* either by dietary restriction-dependent (Y. Zhao et al., 2013) or independent mechanisms (Komura et al., 2013; Nakagawa et al., 2016). The mechanism underlying the dietary effect on the thermotaxis decline does not seem to depend on the activation of the dietary restriction pathway. First, the expression of *pha-4* was low. Second, the lifespan of LAB-fed animals was not necessarily prolonged. Third, *eat-2* mutants, which mimic dietary restriction, did not improve thermotaxis in aged animals fed *E. coli.* Fourth, *kmo-1* and *nkat-1* genes involved in dietary restriction-dependent beneficial effects on associative learning (Vohra, Lemieux, Lin, & Ashrafi, 2017) did not affect the dietary effects on thermotaxis of aged animals. Fifth, no correlation was observed between the thermotaxis ability of aged animals and body size or fat accumulation.

Different LAB activate distinct genetic pathways such as insulin and IGF-1 signaling (IIS) pathway important for lifespan regulation and p38 mitogen-activated protein kinase (MAPK) pathway important for innate immunity. *Lb. rhamnosus* and *B. longum* extend the lifespan of *C. elegans* by modulating the IIS pathway consisting of DAF-2 and DAF-16 (Grompone et al., 2012; Sugawara & Sakamoto, 2018). *B. infantis* extends the lifespan of *C. elegans* via the PMK-1 p38 MAPK pathway and a downstream transcription factor SKN-1, an ortholog of mammalian Nrf, but not via DAF-16 (Komura et al., 2013). The PMK-1 pathway is also activated by *Lb. acidophilus* and *Lactobacillus fermentum* (Y. Kim & Mylonakis, 2012; Park et al., 2018). Animals fed a lactic acid bacterium, *Weissella*, show higher expression of *daf-16*, *aak-2,* and *jnk-1,* and extend lifespan in these genes-dependent manners (Lee et al., 2015). the *daf-16* pathway is also involved in thermotaxis (Kodama et al., 2006; H. Murakami et al., 2005) and salt-avoidance behaviors (Tomioka et al., 2006). In our condition, *daf-16* did not strongly affect thermotaxis at D1, while it was necessary for the maintenance of thermotaxis of *Lb. reuteri*-fed aged animals. This result suggests that DAF-16 had a specific role in the *Lb reuteri*’s effects on aged animals. Since *daf-16* functions in neurons (Fig. 8C) and has neuron-specific targets (Kaletsky et al., 2016), differential expressions of these genes with different diets might affect thermotaxis behavior. Indeed, our transcriptome analysis revealed that differentially expressed genes between animals fed *E. coli* or homofermentative LAB and those fed heterofermentative LAB were enriched with DAF-16 targets. Interestingly, our transcriptome data suggested that DAF-16 was activated in animals fed *E. coli* or homofermentative LAB although heterofermentative *Lb. reuteri*-fed aged animals showed *daf-16* dependency on thermotaxis. The regulation of DAF-16 in specific neurons might be opposite from that in the whole body. Single-cell transcriptome analysis for aged animals fed different diets will address this question in the future (Cao et al., 2017).

### Bacterial screen to address age-dependent phenotypes

Even with *C. elegans*, it is challenging to address age-dependent neuronal phenotypes because powerful forward genetic screens are not readily applicable to aged animals. Our study showed that bacterial screen could be useful for generating phenotypic diversity and addressing underlying molecular mechanisms in aged animals. The bacterial screen has been applied to various *C. elegans* phenotypes. Watson *et al*. carried out unbiased mutant screens of *Escherichia coli* and *Comamonas aquatica* to identify bacterial genes that affect the “dietary sensor” in *C. elegans*, which increases the GFP intensity when fed *Comamonas*; they found that mutations in genes involved in vitamin B12 biosynthesis/import increase *C. elegans* dietary sensor activity (Watson et al., 2014). Zhou et al. screened 13 LAB and found that *Lactobacillus zeae* protects *C. elegans* from enterotoxigenic *E. coli* (M. Zhou et al., 2014). Given that *C. elegans* has its natural microbiota (Berg et al., 2016; Dirksen et al., 2016; Samuel et al., 2016; Zhang et al., 2017), the nervous system of animals in a natural environment may be affected by complex bacteria. Indeed, a recent study has revealed that tyramine produced from commensal bacteria affects *C. elegans* avoidance behavior (O’Donnell et al., 2020). Hence, bacterial screens will provide a unique angle of understanding for *C. elegans* research.

## Materials and Methods

### Worm maintenance and strains

*C. elegans* strains were maintained at 23 °C on Nematode Growth Medium (NGM) plates with *E. coli*, OP50, as previously reported (Brenner, 1974), except CB1370, which was maintained at 15 °C. N2 (Bristol) was used as the wild type. The following mutant strains were used for thermotaxis assays: CB1370 *daf-2(e1370ts)*, DA1116 *eat-2(ad1116),* CF1038 *daf-16(mu86)*, IK0656 *tax-6(db60)*, NUJ69 *kmo-1(tm4529)*, NUJ71 *nkat-1(ok566)*. NUJ69 *kmo-1(tm4529)* is a one-time outcrossed FX04529 *kmo-1(tm4529)* strain. NUJ71 *nkat-1(ok566)* is a two-time outcrossed RB784 *nkat-1(ok566)* strain. The transgenic strains were generated as described below.

### CRISPR knockout

To generate a *daf-16* allele that affects only the exon of the b isoform, we used the Co-CRISPR strategy (H. Kim et al., 2014). tracrRNA, *daf-16* crRNA (5’-GUCAUGCCAGAUGAAGAACA-3’), and *dpy-10* crRNA (5’-GCUACCAUAGGCACCACGAG-3’) were annealed and injected into N2 animals with *eft-3*p::Cas9::NLS-3’UTR(*tbb-2*) plasmids. After picking dumpy and/or roller F_1_ animals on individual plates, the mutation of the *daf-16* locus was detected by PCR and confirmed using Sanger sequencing. We obtained *knj36* which introduced 7-bp deletion in the first exon of the *daf-16* b isoform, corresponding to the intron of other isoforms. NUJ298 *daf-16(knj36) I* was used for thermotaxis assays.

### Plasmids and single-copy insertion of transgenes

Single-copy insertions of transgenes were generated using Cas9-based homologous recombination following the established protocol (Andrusiak et al., 2019). *daf-16*bp::*daf-16*b (gDNA) was cloned into the pCR8 vector (Thermo Fisher Scientific) vector using Gibson Assembly (New England Biolabs) (pKEN891). pKEN891 was recombined with the pCZGY2729 repair template vector using the LR reaction (Gateway Technology, Thermo Fisher Scientific) to generate the pKEN1016 repair template vector. pKEN1016 contains homology arms for *cxTi10882* site on chromosome IV and *rps-0*p::Hyg^R^ in addition to the *daf-16*b promoter (4.9 kbp) and *daf-16*b gDNA. To swap promoters of pKEN1016, XhoI and AgeI sites were introduced before and after the *daf-16*b promoter, respectively (pKEN973). pKEN973 was digested with XhoI and AgeI, and the promoter was substituted to *myo-2*p (1.0 kbp, pKEN976), *myo-3*p (2.4 kbp, pKEN975), or *rgef-1*p (3.5 kbp, pKEN974) using Gibson assembly.

We injected the repair template described above and *eft-3*p::Cas9 + *cxTi10882* sgRNA (pCZGY2750) into N2 or CF1038 *daf-16(mu86)* with three red markers originally used for MosSCI insertion (Frøkjær-Jensen et al., 2008): *rab-3*p::mCherry (pGH8), *myo-2*p::mCherry(pCFJ90), and *myo-3*p::mCherry (pCFJ104). The animals with the single copy insertion were selected based on the hygromycin-resistance (Hyg^R^) and the absence of red fluorescence. The following strains were used for the rescue experiments of thermotaxis assays: NUJ306 *daf-16(mu86) I; knjSi17[daf-16bp::daf-16b] IV*, *NUJ368 daf-16(mu86) I; knjSi19[myo-2p::daf-16b] IV*, NUJ373 *daf-16(mu86) I; knjSi18[myo-3p::daf-16b] IV*, NUJ372 *daf-16(mu86) I; knjSi22[rgef-1p::daf-16b] IV*.

### Bacterial plates

*E. coli*, OP50 or HT115, was inoculated into Super Broth (32 g Bacto Tryptone (BD), 20 g Bacto Yeast extract (BD), 5 g NaCl (Wako), 5 mL 1M NaOH in 1 L water) and cultured overnight at 37 °C. LAB strains, provided by Megmilk Snow Brand company (Table S1), were inoculated into the liquid medium from glycerol stocks and cultured in the conditions described in Supplementary Table 1. Bacterial cells were collected by centrifugation at 7,000x g for 10 min at 4 °C. Cells were washed twice with sterile 0.9% NaCl solution. The washed bacteria were adjusted to a final concentration of 0.1 g/ml (wet weight) in NG buffer (25 mM K-PO_4_ (pH6), 50 mM NaCl, 1 mM CaCl_2_, 1 mM MgSO_4_). For heat killing, 0.1 g/ml bacteria in tubes were incubated for 1 h in a 65 °C incubator or 10 min in boiled water. By this treatment, bacterial colony-forming unit (cfu) became <1.0×10^2^ cfu/ml, which is at least 10^8^ lower than live bacteria (>1.5×10^10^). For the mixed condition, bacteria were mixed to make the final concentration of 0.1 g/ml in total before spread on NGM plates. To crush bacteria, bacterial suspension was vigorously vibrated with glass beads at 4,200 rpm for 15 cycles of 30-sec ON and 30-sec OFF using a bead-based homogenizer (MS-100R, Tomy). Two hundred microliters of the bacterial suspension were spread onto 60-mm NGM plates and dried overnight. NGM plates with peptone were used except for thermotaxis to see the effect of peptone and lifespan assays, in which NGM plates without peptone were used.

### Preparation of aged animals fed different bacteria

For behavioral assays, synchronized eggs were prepared by bleaching gravid hermaphrodites using 0.5x household bleach in 0.5 M NaOH and placed onto NGM plates with OP50. The eggs were cultivated at 23 °C for 72 hours to obtain day one adults (D1). For the thermotaxis of *daf-2* mutants, eggs of N2 and CB1370 *daf-2(e1370)* were incubated at 15 °C for 96 hours and subsequently at 23 °C for 24 hours to obtain D1. D1 animals were washed with NG buffer and transferred to NGM plates with OP50 or LAB every day for thermotaxis of aged animals. To expose animals to the bacterial odor, NG buffer as control or 0.1 g/ml bacterial solution was spotted on the lid of bacterial plates described above. Animals were cultivated on the upside-down plates as described in Figure 5D. For the thrashing and locomotion assay, animals were transferred individually by picking instead of washing.

### Thermotaxis assay

Animals were cultivated at 23 °C (T_cult_=23 °C) unless otherwise noted. Population thermotaxis assays with a linear thermal gradient were performed as described (Ito et al., 2006). Fifty to 250 animals on cultivation plates were washed with M9 and placed at the center of the assay plates without food and with a temperature gradient of 17-23 or 20-26 °C. The temperature gradient was measured to be ∼0.5 °C/cm. After letting them move for 1 h, the animals were killed by chloroform. The number of adult animals in each of eight sections along the temperature gradient (Fig. 1A) was scored under a stereomicroscope. The fraction of animals in each section was plotted on histograms. The performance and thermotaxis indices were calculated, as shown in Fig. 1A and Fig. 3C, respectively. For temperature shift assays, T_cult_ was shifted 24 h prior to the assay.

### Thrashing assay

A thrashing assay was performed, as previously described with a few modifications (Tsalik & Hobert, 2003). Animals were washed with NG buffer and transferred with a drop of NG buffer onto an NGM plate without food using a capillary glass pipet. In liquid, animals show lateral swimming movements (thrashes). We defined a single thrash as a complete movement through the midpoint and back and counted the number of thrashes for 30 seconds.

### Motility assay

Assay plates were prepared by placing circular filter paper with a one-inch hole on NGM plates with OP50 or LAB and soaking the paper with ∼100 µl of 100 mM CuCl_2_. A single animal was transferred to an assay plate with the cultured bacteria and left at 23 °C for three minutes. The images of the bacterial lawn were captured by a digital camera (Fujifilm) through an eyepiece of a stereomicroscope, Stemi 508 (Zeiss). The trajectory of an animal on the lawn was traced using FIJI (Schindelin et al., 2012) and measured as the distance of locomotion.

### Lifespan assay

Animals were synchronized by bleaching gravid adults and grown with regular NGM plates with OP50 until D1. D1 animals were washed three times with M9 buffer and transferred to peptone-free NGM plates supplemented with 50 mg/ml OP50 or LAB. Animals were transferred to new plates every day until they became D4 and every other day afterward. Dead animals were defined as having no voluntary movement after several touches on the head and tail and counted every day. Four independent sessions with 25 animals per session were combined for each condition.

### Phylogenetic tree

16S rRNA sequences of LAB were obtained from the Genome database of NCBI (http://www.ncbi.nlm.nih.gov/genome/). The accession numbers are shown in Table S1. The phylogenetic tree was inferred by the Neighbor-Joining method based on the 16S rRNA gene sequence of model LAB strains. The evolutionary distances were computed using the Maximum Composite Likelihood method conducted in MEGA X (Hall, 2013).

### RNA sequencing

Non-gravid young adult animals were used to avoid the effect of eggs inside the body. D5 animals fed *E. coli* or LAB (*Lb. gasseri*; *Lb. delbrueckii*, *P. pentosaceus*, *Lb. reuteri*, *Lb. rhamnosus*, and *Lb. plantarum)* were prepared as described above. Total RNA was extracted from whole animals using RNAiso Plus reagent (Takara) and sequenced using NovaSeq 6000 System by Macrogen Corp. Japan. We detected 13329 genes. The fragments per kilobase of exon per million mapped fragments (FPKM) of D1, D5 (*E. coli*), and D5 (LAB, 6 in total) were used to perform the principal component analysis (PCA) based on the variance-covariance matrix using R program. The eigenvalues of each transcript were ranked by the percentage of explained variance for each principal component, and the top and bottom 5% (666 genes each) were used to perform gene ontology (GO) analysis using Metascape (Y. Zhou et al., 2019).

To further analyze the difference between D5 animals fed *E. coli* or homofermentative LAB (*Lb. gasseri*; *Lb. delbrueckii*) and D5 animals fed heterofermentative LAB (*P. pentosaceus*, *Lb. reuteri*, *Lb. rhamnosus*, and *Lb. plantarum*), we compared the expression of genes between these two groups and extracted differentially expressed genes as those with p-value<0.05 by Student’s t-test. Those differentially expressed genes were used to perform GO analysis using Metascape and compared with DAF-16-regulated genes (Tepper et al., 2013). Gene enrichment was calculated using Gene Set Enrichment Analysis (GSEA) (Subramanian et al., 2005).

### Statistical analyses

Box-and-whisker plots represent medians as center lines; boxes as first and third quartiles; whiskers as maximum and minimum values except for outliers, which are 1.5 times greater than the upper limit or 1.5 times smaller than the lower limit of the interquartile range; dots as outliers. In some figures, all the data points were overlayed on the box-and-whisker plots. We used Student’s t-test to compare two samples and one-way or two-way ANOVA, followed by Dunnett’s or Tukey-Kramer test to compare multiple samples using R (R core team, https://www.R-project.org/, Vienna, Austria) or GraphPad Prism 7.0 (GraphPad Software, La Jolla, CA). In all figures, *p < 0.05, **p < 0.01, ***p<0.001. p >0.05 is considered as not significant (ns).

## Acknowledgments

Megmilk Snow Brand Company supported this work. We thank members of the Nutritional Neuroscience laboratory and the Mori laboratory for their comments on the manuscript. Wei Huang and Pauline Rouillard helped with basic experiments. *C. elegans* mutant strains were provided by Caenorhabditis Genetics Center (CGC), funded by the NIH Office of Research Infrastructure Programs (P40 OD010440), and Dr. Shoehei Mitani of the National Bioresource Project of Japan. Dr. Yishi Jin provided pCZGY2729 and pCZGY2750 plasmids.

## Supplementary information

### Supplementary Materials and Methods

#### AFD and AIY imaging

To examine if AFD and AIY experience cell death at D5, NUJ296 *knjIs15[gcy-8Mp::GCaMP6m + gcy-8Mp::tagRFP + ges-1p::tagRFP]* and IK1144 *njIs26[AIYp::GCaMP3 + AIYp::tagRFP + ges-1p::tagRFP]* were imaged, respectively. D1 and D5 animals were immobilized using 1 mM levamisole in M9, mounted on an agarose pad, and imaged using an Axio Imager.A2 equipped with a Plan-Apochromat 63x/1.4 oil objective (Zeiss). TagRFP signals in the cell body of AFD or AIY were visualized by green LED (555/30 nm) of Colibri 7 light source, and a quad-band path filter ser 90 HE LED (Zeiss) and used to evaluate possible cell death. We did not observe any loss of cell bodies in young or aged animals.

#### Food recognition assay

The food recognition assay was performed as previously described with a few modifications (Sawin et al., 2000). Assay plates with food were prepared by spreading OP50 onto NGM plates. For well-fed animals, animals were washed twice in S basal buffer (Brenner, 1974) and transferred using a capillary glass pipette into a drop of the buffer on an assay plate with or without food. Five minutes after transfer, the number of body bends in 20 s intervals was counted. For starved animals, 5–15 animals were washed twice in S basal buffer and incubated on NGM plates without food for 30 min. After transferring them on assay plates with or without food, we measured the number of body bends.

#### Salt avoidance assay

For a gradient assay of salt chemotaxis (Saeki et al., 2001), a salt gradient was formed overnight by placing an agar plug containing 100 mM of NaCl (5 mm diameter) 2 cm away from the edge of the 90 mm assay plate (2% Bacto Agar, 5 mM K-PO_4_ (pH 6.0), 1 mM CaCl_2_, 1 mM MgSO_4_). D1, *E. coli*-fed D5, and *Lb. reuteri*-fed D5 animals were divided into three groups. The first group of animals received no conditioning (Naïve). For conditioning, animals were washed three times with chemotaxis buffer (5 mM KPO_4_ (pH 6.0), 1 mM CaCl_2_, 1 mM MgSO_4_), transferred to the same buffer with 20 mM NaCl (NaCl-conditioned) or without NaCl (Mock-conditioned), and incubated at 25°C for 1 hr. These animals were placed at the center of the assay plates and then incubated at 25°C for 30 min. The chemotaxis index was calculated as (N_NaCl_ − N_control_)/(N_total_− N_origin_) as indicated in Figure S4B. One hundred to two hundred animals were used in each assay.

For a quadrant assay of salt taxis (Wicks et al., 2000), we used a compartmentalized plate (Falcon X, Becton Dickinson Labware) as an assay plate (2% Bacto Agar, 5 mM K-PO_4_ (pH 6.0), 1 mM CaCl_2_, 1 mM MgSO_4_). The plates were freshly prepared on the day of the assay with two different agar solutions, 0 mM NaCl and 25 mM NaCl. D1, *E. coli*-fed D5, and *Lb. reuteri*-fed D5 animals were divided into three groups. The first group of animals received no conditioning (Naïve). For conditioning, animals were washed three times with chemotaxis buffer (5 mM K-PO_4_ (pH 6.0), 1 mM CaCl_2_, 1 mM MgSO_4_), transferred to the same buffer with 100 mM NaCl (NaCl-conditioned) or without NaCl (Mock-conditioned), and incubated at room temperature for 15 min. These animals were placed at the center of the assay plates and then incubated at room temperature for 10 min. The chemotaxis index was calculated as (N_NaCl_ − N_control_)/(N_total_− N_origin_) as indicated in Figure S4B. One hundred to two hundred animals were used in each assay.

#### Gram staining

Bacteria are fixed with methanol and stained using Gram Color Kit (Muto Pure Chemicals Co., Ltd., Tokyo, Japan). Stained bacteria are imaged using an Axio Imager.A2 equipped with a Plan-Apochromat 63x/1.4 oil objective (Zeiss).

#### FITC-labeling of bacteria

To examine whether *C. elegans* ingests bacteria, we used fluorescently labeled bacteria. Bacterial cells were incubated with Phosphate-Buffered Saline (PBS, Takara) (unlabeled) or 0.1 mg/ml FITC-I (Wako) in PBS for one hour, washed with PBS three times, and resuspended with PBS at 0.1 g/ml. For the mixed condition, an equal amount of unlabeled and FITC-labeled bacteria were mixed. Bacteria were spread on NGM plates and dried. D1 adult animals were placed on the bacterial plate and incubated at 23 °C for 20 min. Excess amounts of fluorescent bacteria were removed by letting worms crawl on an NGM plate without food for a few minutes. Animals were imaged after washing using an Axio Imager.A2 equipped with a Plan-Apochromat 63x/1.4 oil objective (Zeiss).

#### Quantitative RT-PCR

RNA was prepared as described in the RNA sequencing section. Two micrograms of total RNA were reverse transcribed to cDNA with a mixture of random and oligo dT primers using ReverTra Ace qPCR RT Master Mix with gDNA Remover (TOYOBO). The cDNA and gene-specific primers were used for qPCR reaction with THUNDERBIRD SYBR qPCR Mix (TOYOBO), and the products were detected using a LightCycler 96 System (Roche). The gene-specific primers were used for *pha-4*: KN1370, 5’-GGTTGCCAGGTCCCCTGACA-3’; KN1371, 5’-GCCTACGGAGGTAGCATCCA-3’. *cdc-42* was used as a reference because it is stable and unaltered during aging (Hoogewijs, Houthoofd, Matthijssens, Vandesompele, & Vanfleteren, 2008; Mann, Van Nostrand, Friedland, Liu, & Kim, 2016) (KN1170, 5’-CTGCTGGACAGGAAGATTACG-3’; KN1171, 5’-CTCGGACATTCTCGAATGAAG-3’).

## Supplementary Figure Legends

**Figure S1.**
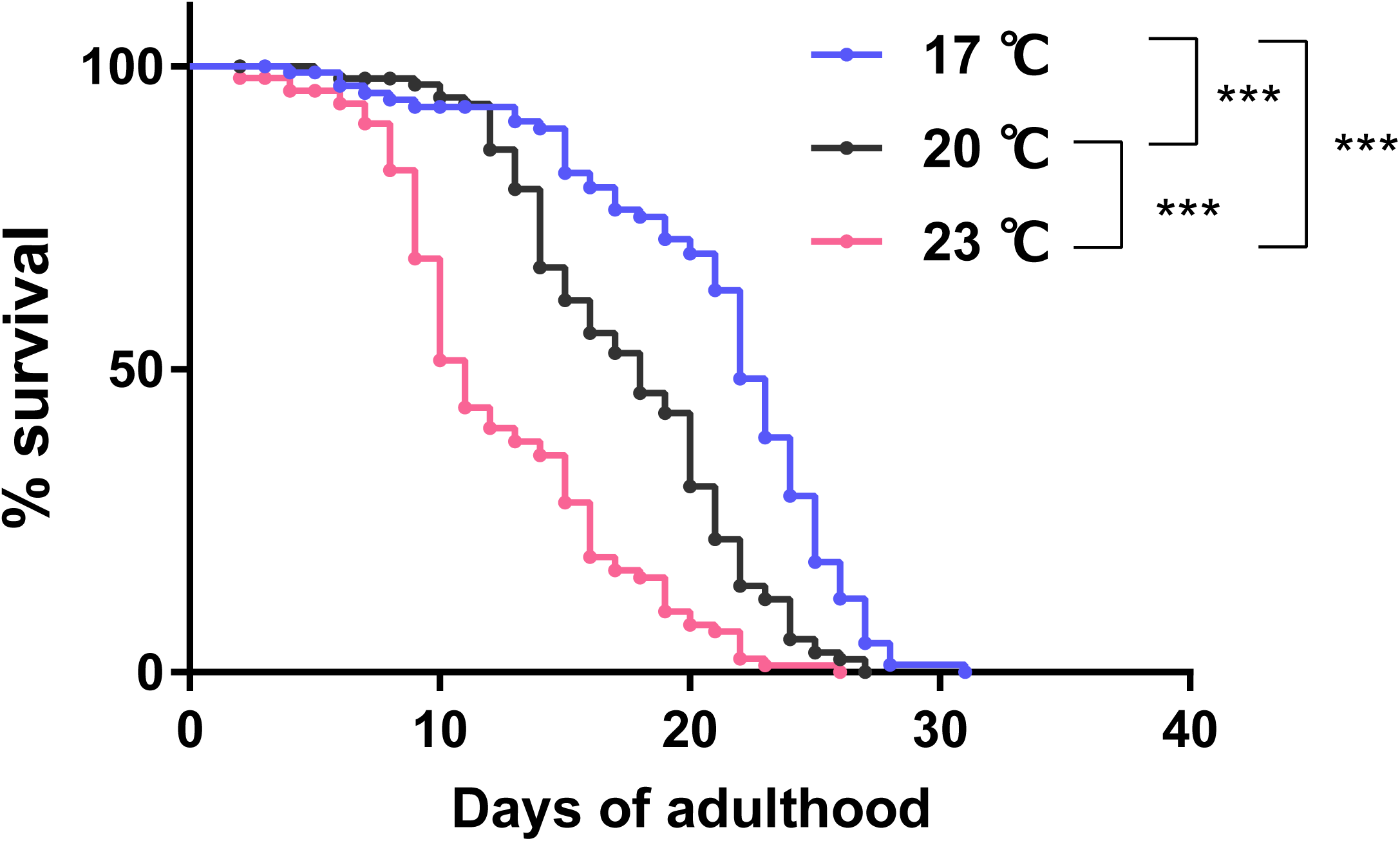
Survival curve of animals cultivated at different temperature. Survival curves of wild-type animals cultivated with *E. coli* from eggs at the indicated temperatures. N=4 experiments with 25 animals/experiment (100 animals in total). Statistics: Log-rank test, p***<0.001.

**Figure S2.**
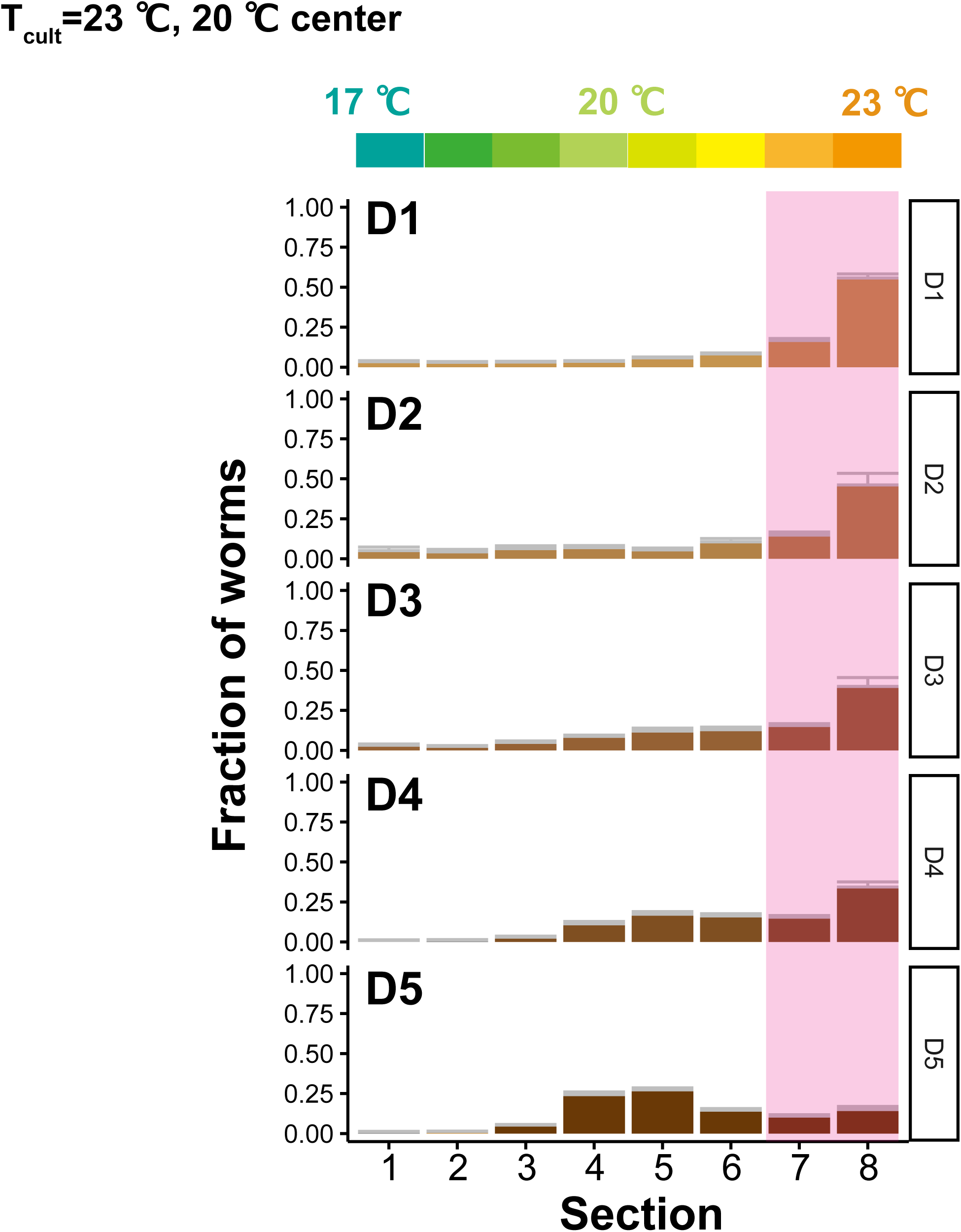
Distributions of animals on the temperature gradient at different age. The distributions of animals at D1 - D5 on the temperature gradient after thermotaxis assays. Animals were cultivated at 23 °C and placed at the center of the 17-23 °C gradient. Pink rectangles indicate the sections around the T_cult_.

**Figure S3.**
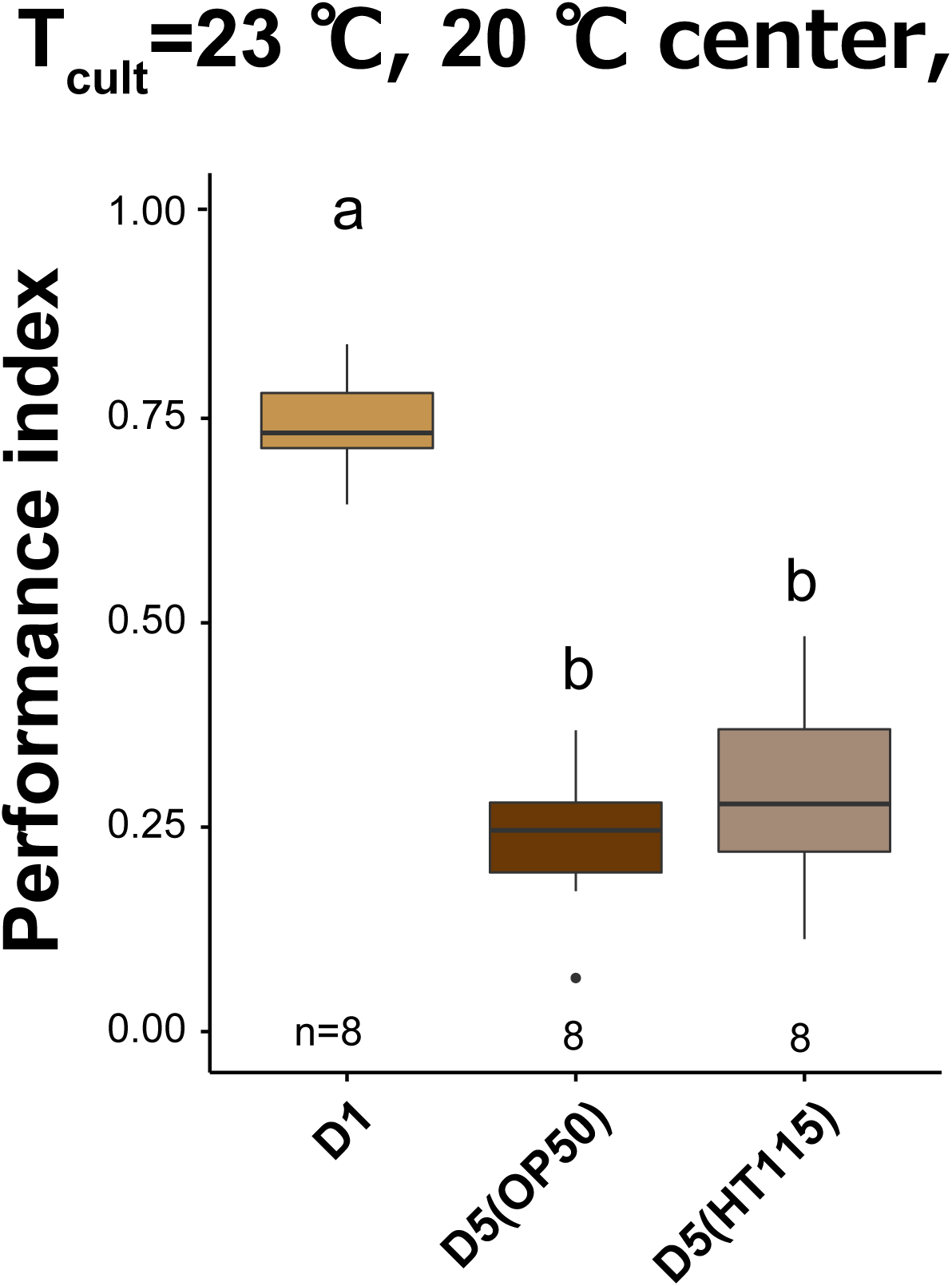
HT115-fed animals showed thermotaxis decline. Box plots summarizing thermotaxis performance indices. Animals were cultivated with OP50 until D1 and then with OP50 or HT115 until D5. The number of experiments is shown. Statistics: The mean indices marked with distinct alphabets are significantly different (p < 0.05) according to One-way ANOVA followed by Tukey–Kramer test.

**Figure S4.**
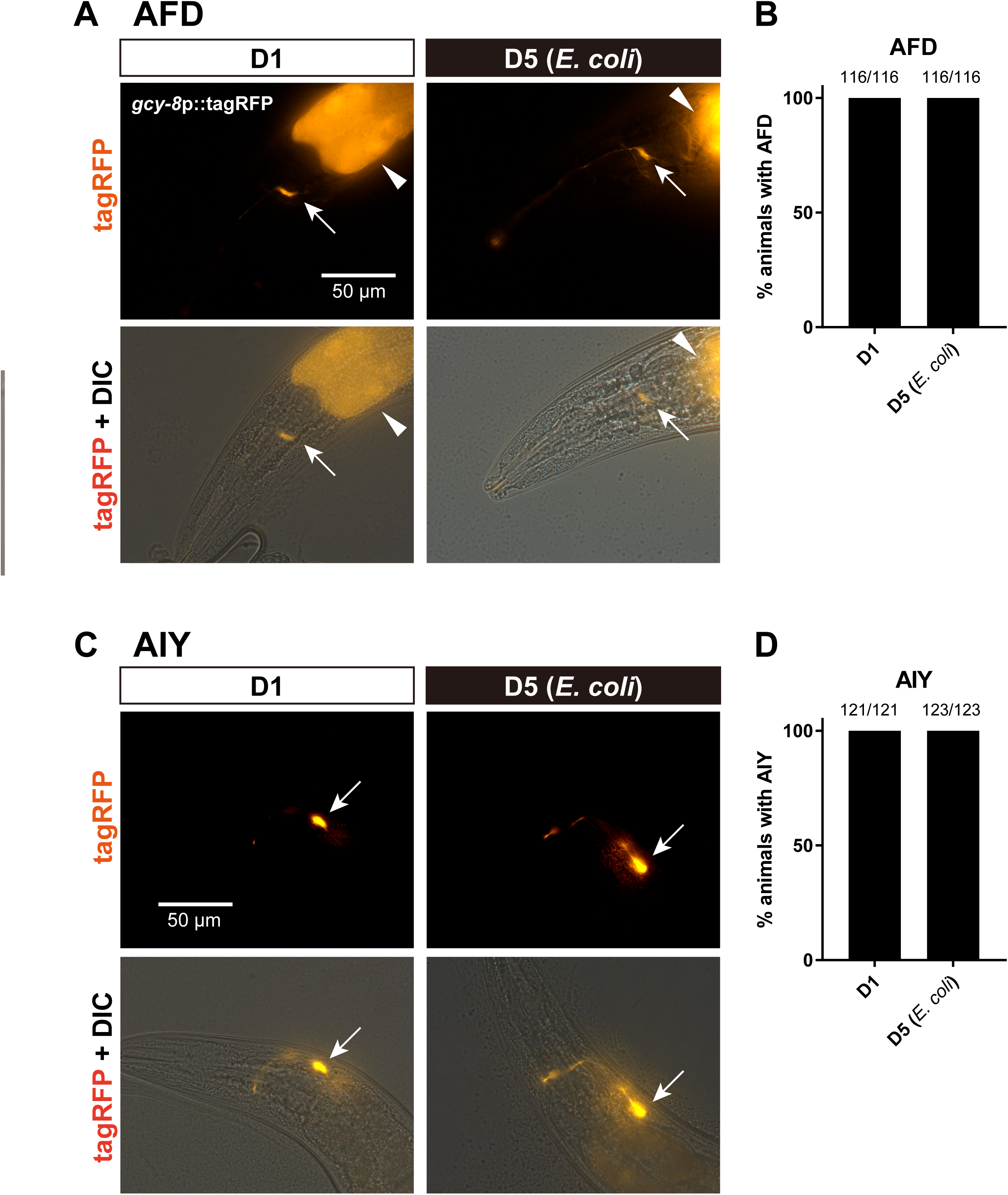
AFD and AIY are alive at D5. TagRFP of NUJ296 *knjIs15[gcy-8Mp::GCaMP6m + gcy-8Mp::tagRFP + ges-1p::tagRFP]* and IK1144 *njIs26[AIYp::GCaMP3 + AIYp::tagRFP + ges-1p::tagRFP]* were imaged to visualize AFD and AIY, respectively. (A and C) Representative images of the head of C. elegans for tagRFP (top panels) or tagRFP + Differential interference contrast (DIC) (bottom panels). Scale bar: 50 µm. Arrows indicate the cell bodies of the neurons. Arrowheads in (A) indicate tagRFP expression in the intestine used as a co-injection marker. (B and D) Bar graphs show the percentage of animals with AFD (B) and AIY (D) in D1 and D5, based on tagRFP signals in the cell body. The number indicates the number of animals with visible neurons over the total number of animals examined.

**Figure S5.**
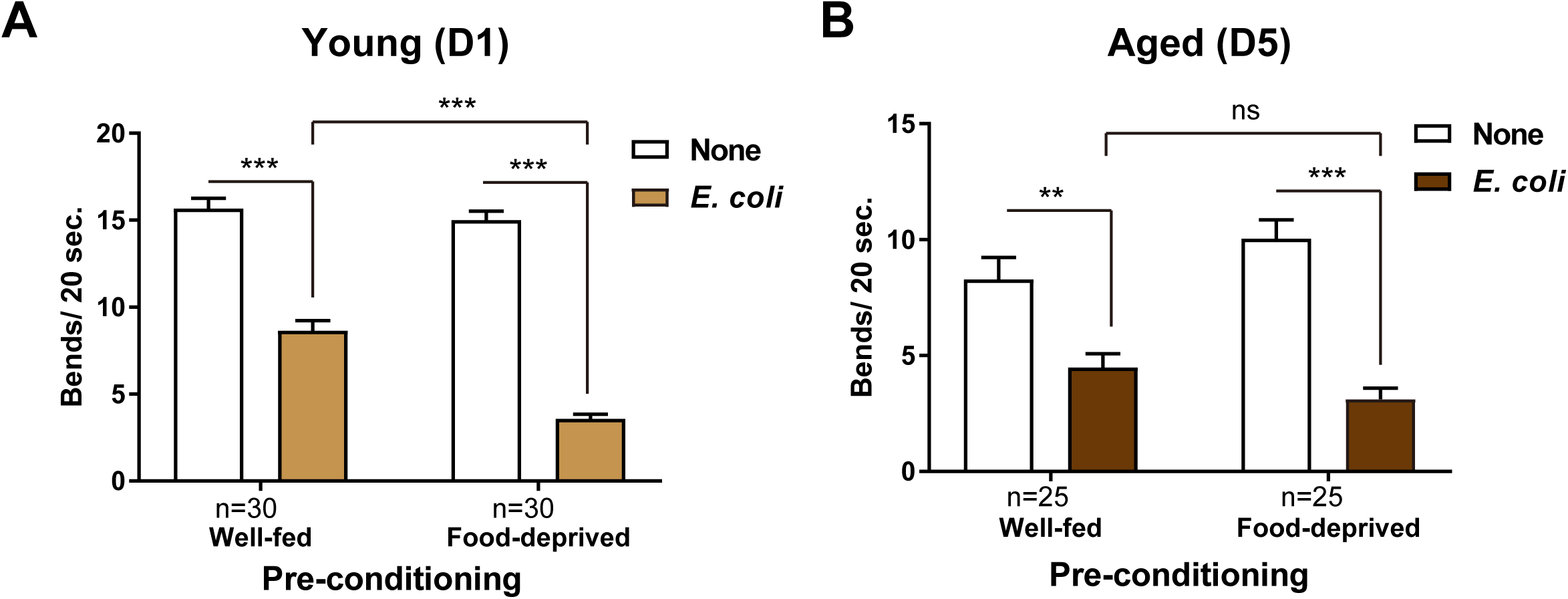
*E. coli*-fed aged animals sense food. (A and B) Food recognition assays of D1 in (A) and D5 adults in (B). Animals were pre-conditioned with (well-fed) or without (food-deprived) *E. coli* and assayed on plates with or without *E. coli*. Animals’ locomotion was evaluated by body bends in 20 sec. The presence of food on the assay plate slows down the locomotion of well-fed animals (basal slowing response). Pre-conditioning animals without food enhanced the basal slowing response (enhanced slowing response). The number of animals examined is shown. Error bars: S.E.M. Statistics: Two-way ANOVA with Turkey’s multiple comparison test, ***p<0.001; **p<0.01; ns, p>0.05.

**Figure S6.**
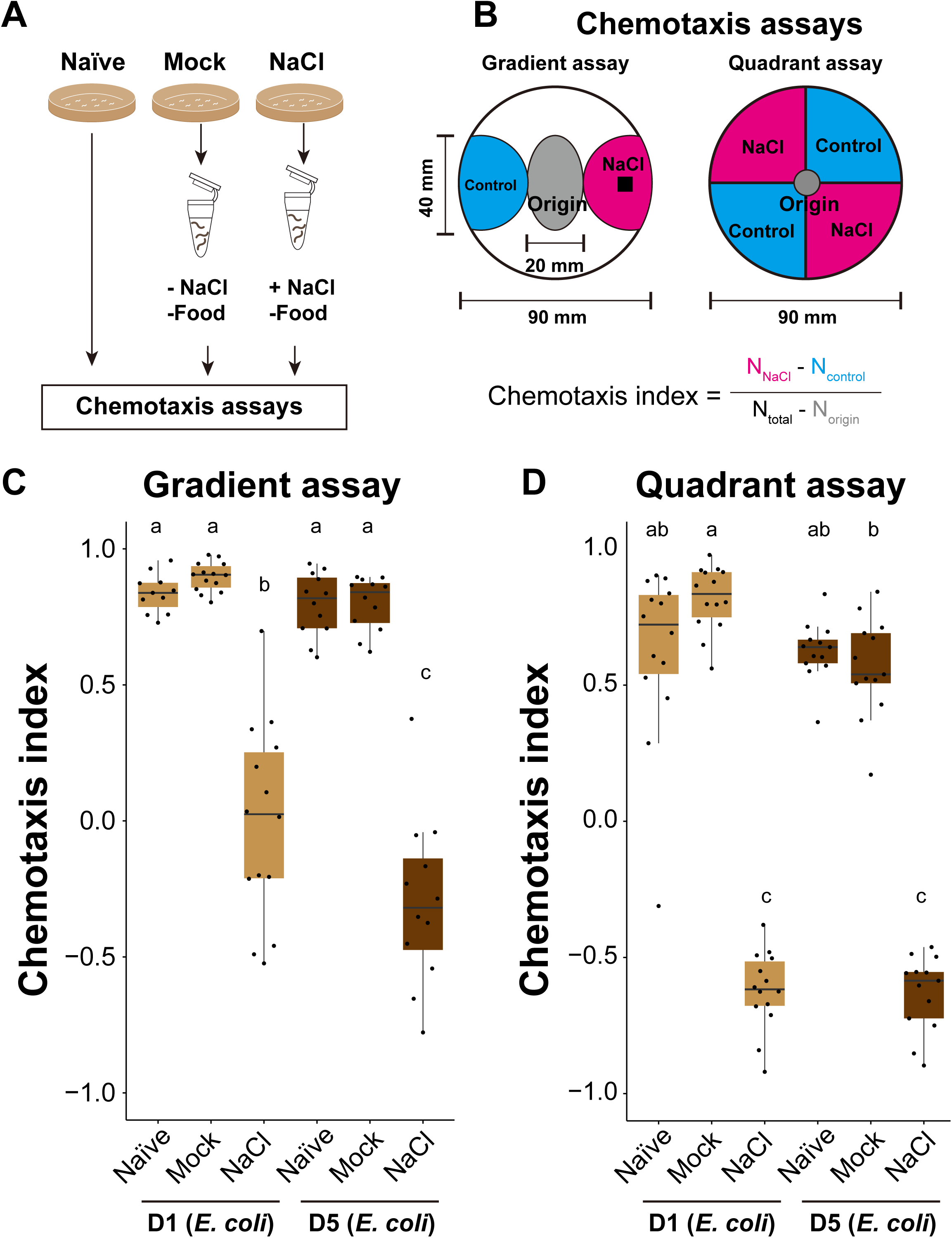
Aged *E. coli*-fed animals perform salt avoidance behavior irrespective of diet. (A) Schematic of conditioning of animals. Animals were conditioned in the absence of food with (NaCl) or without (Mock) NaCl. (B) Schematic of salt avoidance assays. Gradient and quadrant assays were used to evaluate salt avoidance behavior. The chemotaxis index was calculated using the indicated formula based on the number of animals in each area after the assay. (C and D) Box plots summarizing chemotaxis indices of salt avoidance assays. The number of experiments is shown. One-way ANOVA followed by Dunnett’s multiple comparison test compared to Naïve condition in each condition, ***p<0.001; ns, p>0.05.

**Figure S7.**
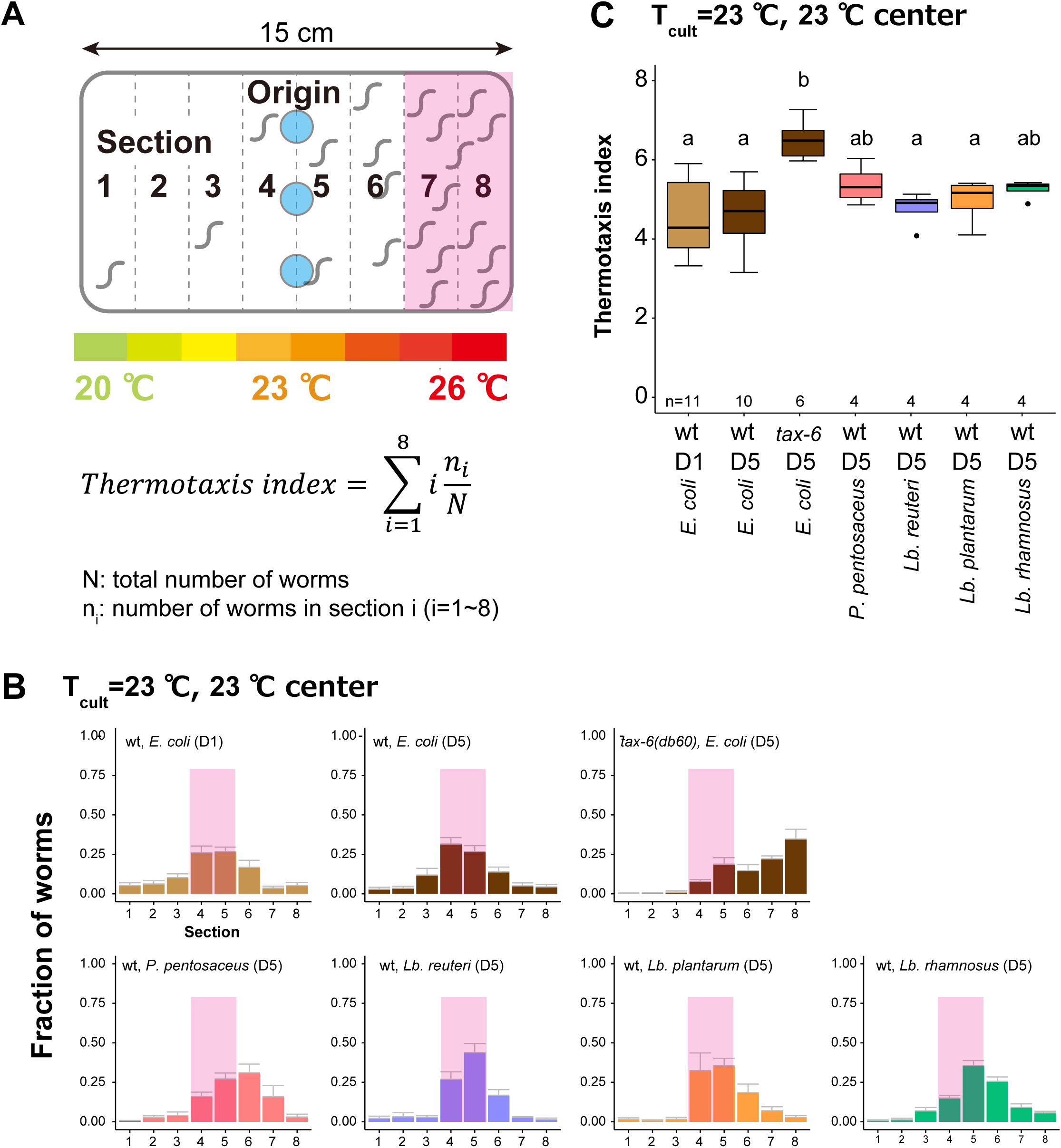
Animals fed heterofermentative LAB are not thermophilic. Animals were cultivated at 23 °C and placed at the center of a 20-26 °C gradient to examine the thermophilicity. D5 animals with indicated genotypes were cultivated with *E. coli* or *Lb. reuteri* from D1. *tax-6(db60)* was used as a thermophilic mutant. (A) Schematic of thermotaxis assay and formula to calculate thermotaxis index. (B) Distribution of animals on the temperature gradient after thermotaxis assays. Pink rectangles indicate the sections around the T_cult_. (C) Box plots summarizing thermotaxis indices of animals under indicated conditions. The number of experiments is shown. Statistics: The mean indices marked with distinct alphabets are significantly different (p < 0.05) according to One-way ANOVA followed by Tukey–Kramer test.

**Figure S8.**
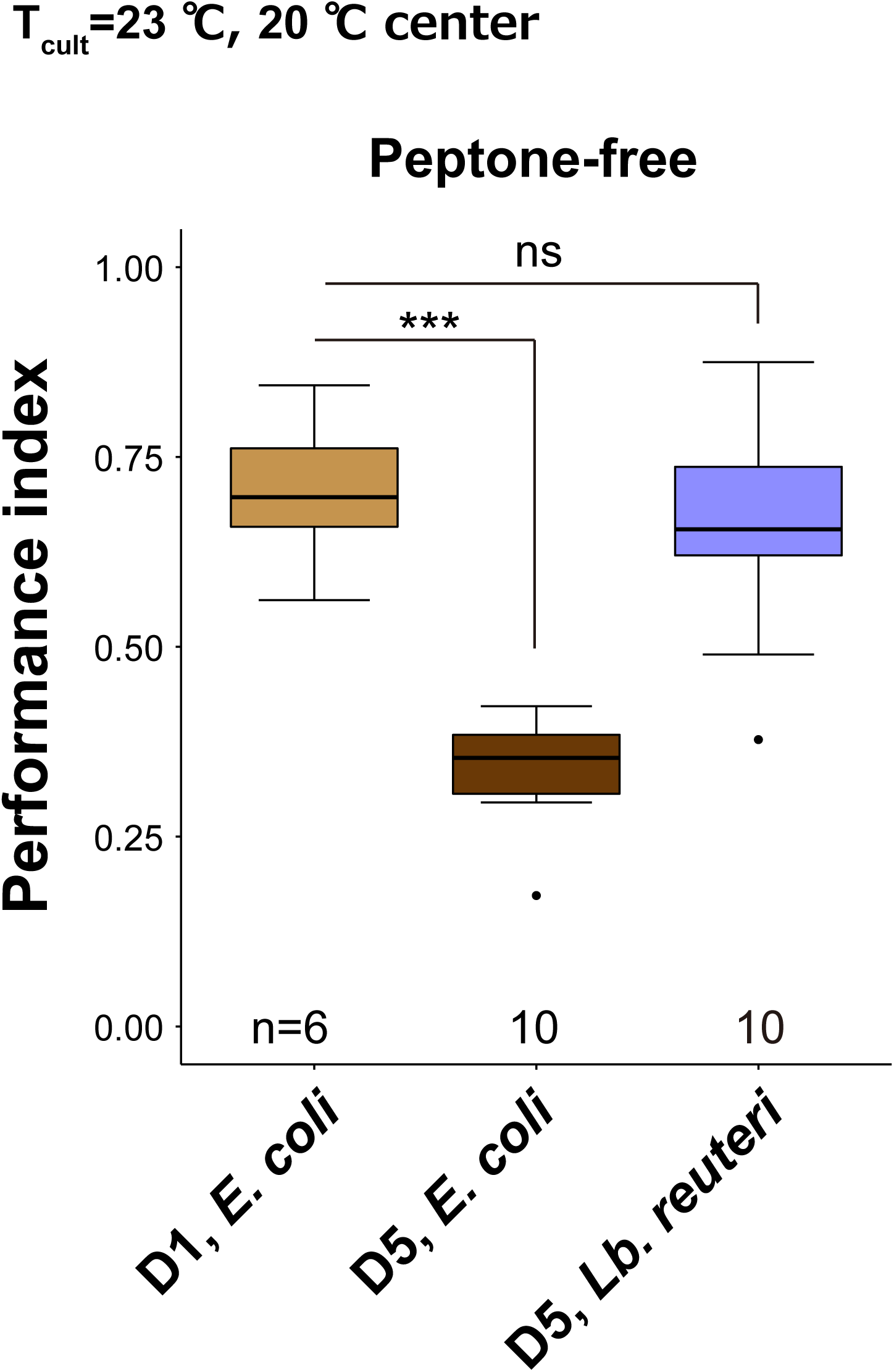
Diets affect thermotaxis of aged animals on peptone-free plates. Box plots summarizing performance indices of animals cultivated on peptone-free NGM plates. D5 animals were cultivated at 23 °C with *E. coli* or *Lb. reuteri* from D1. The number of experiments is shown. Statistics: One-way ANOVA followed by Dunnett’s multiple comparison test compared to the D1 control, p***<0.001; ns, p>0.05.

**Figure S9.**
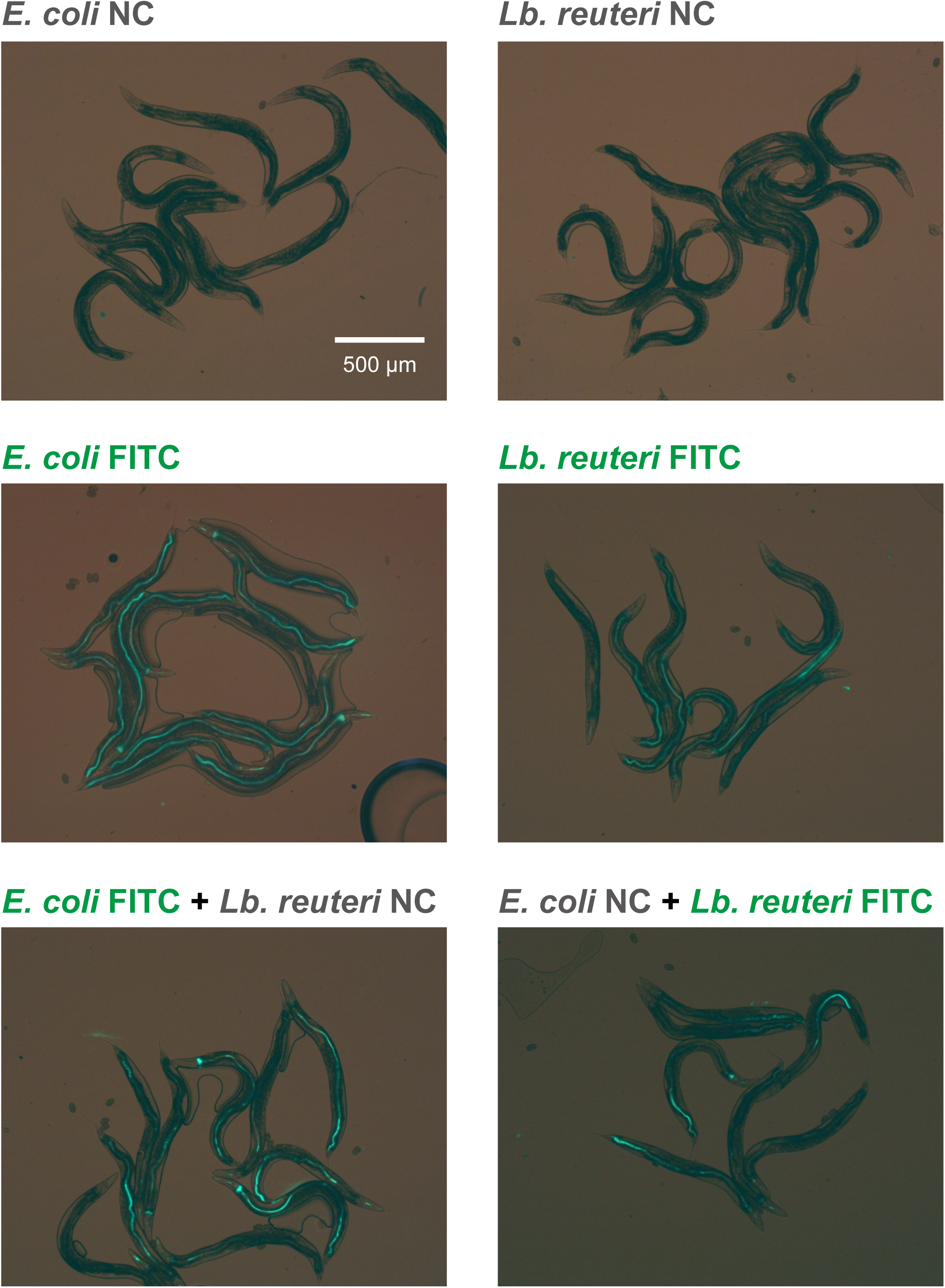
Animals ingest *E. coli* and *Lb. reuteri*. Representative images show fluorescently labeled bacteria in the intestine of *C. elegans*. Animals were cultivated with FICT-labeled *E. coli*, *Lb. reuteri*, or the mixture of fluorescently labeled and unlabeled bacteria. Both fluorescence and transmission light was simultaneously imaged to show that *C. elegans* ingested bacteria.

**Figure S10.**
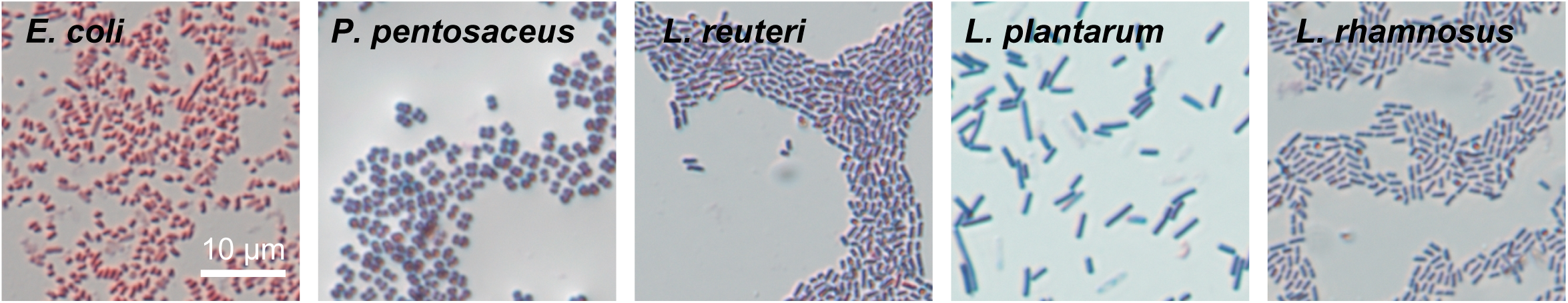
Images of Gram-stained bacteria. Images of Gram-stained *E. coli* and select LAB. *E. coli* and LAB are Gram-negative and positive, respectively. Scale bar: 10 µm.

**Figure S11.**
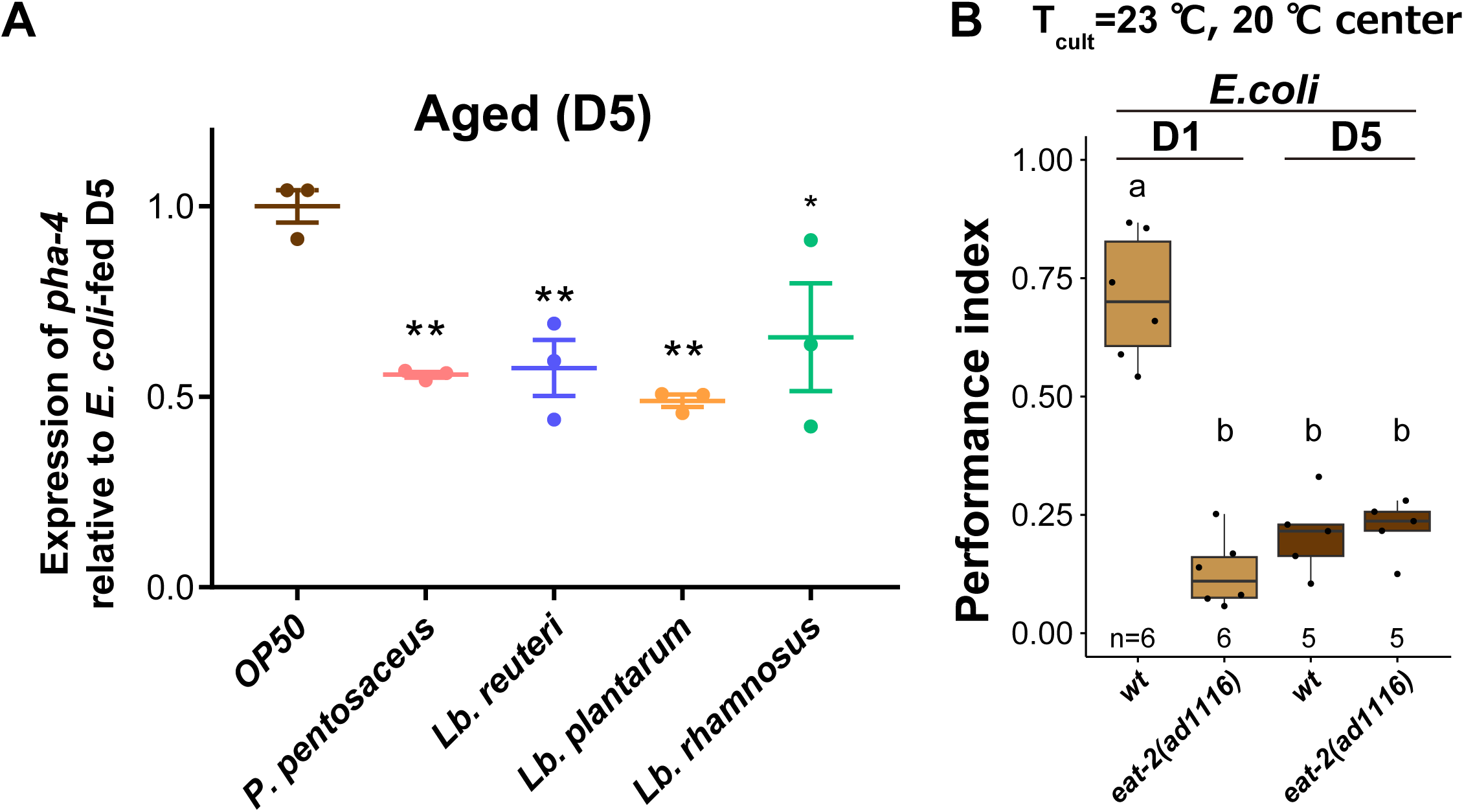
LAB-fed bacteria are not dietary-restricted. (A) Expression of *pha-4* transcripts in aged animals fed indicated LAB relative to aged animals fed *E. coli*. (B) Box plots summarizing thermotaxis indices of wild type and *eat-2* mutant animals cultivated at 23 °C with *E. coli*. The number of experiments is shown. Statistics: Student’s t-test, ns, p>0.05.

**Figure S12.**
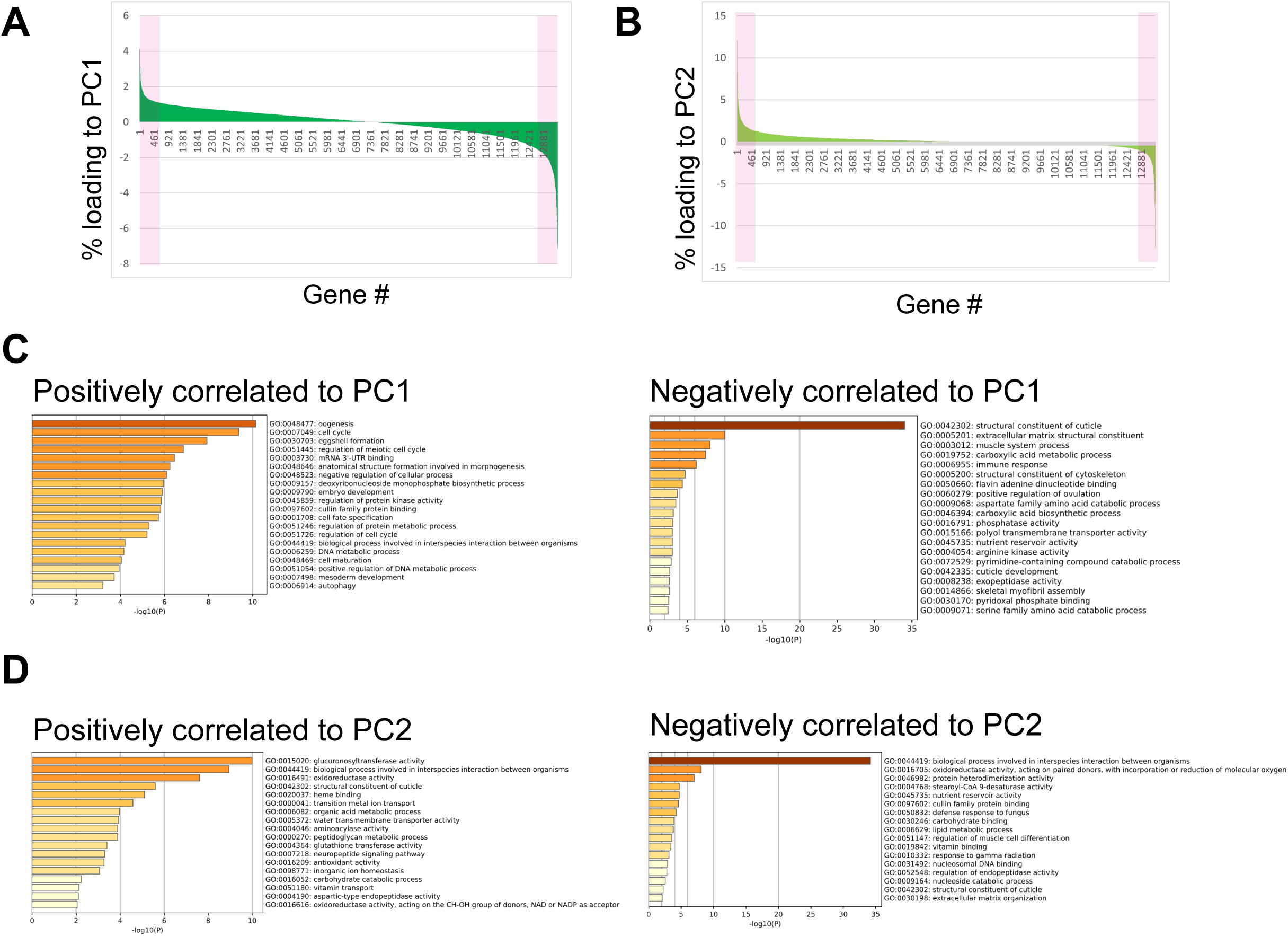
Principal component analysis of the transcriptome data of the animals of different ages and diets. Principal component analysis was carried out for the transcriptome data of D1, and D5 animals fed *E. coli* or LAB (*Lb. gasseri*; *Lb. delbrueckii*, *P. pentosaceus*, *Lb. reuteri*, *Lb. rhamnosus*, and *Lb. plantarum*). Loadings of genes were ranked. (A and B) Loadings of each gene to PC1 (A) and PC2. Magenta indicates the top 5% of genes most correlated and anti-correlated to PC1 or PC2. the x-axis indicates genes, but the orders are different between (A) and (B). (C and D) Gene ontology analysis of the top 5% of genes correlated or anti-correlated with PC1 (C) or PC2 (D).

**Table S1 List of LAB strains**

